# Coupling cell differentiation to dewetting can explain villus elongation

**DOI:** 10.64898/2026.05.14.725076

**Authors:** Dominic K Devlin, Shuji Ishihara, Austen RD Ganley, Nobuto Takeuchi

## Abstract

During vertebrate development, the flat surface of the gut epithelium undergoes a dramatic transformation into densely packed arrays of finger-like projections called intestinal villi. Recent studies show that the villus formation relies on a tissue dewetting process, in which mesenchymal tissues buckle the overlying epithelial layer into periodic folds. However, the mechanisms driving subsequent elongation of these folds into finger-like villi remain largely unexplored. Here, we propose a simple mechanism for villus elongation that couples tissue dewetting to cell differentiation, which emerged as a repeated outcome of multiple independent simulations of an evolutionary-developmental Cellular Potts Model. In this mechanism, a liquid-like mesenchymal tissue continuously differentiates into a solid-like mesenchymal tissue at the interface between them. This differentiation drives the liquid-like tissue to continuously retract from the solid-like tissue in the opposite direction of the interface through dewetting, ultimately creating a finger-like projection. A merit of our proposed mechanism is that it only requires two tissues with different viscosities, high surface tension, and cell differentiation. We develop a simplified phase-field model to determine exactly how villus morphology depends on these three requirements. Since these requirements are satisfied not only in intestinal villi but also in many other developing tissues, we propose that the same mechanism could also drive the elongation of other tissues.

## 1 Introduction

The transformation of cell sheets or spheres into elongated structures is a fundamental process in embryonic and organ morphogenesis [1]. Tissues can achieve this elongation through (i) autonomous mechanisms, or (ii) mechanisms involving the cooperation of neighbouring tissues. Autonomous mechanisms include convergent extension, in which a tissue narrows and lengthens along perpendicular axes (e.g., the embryonic notochord [2]), and oriented cell divisions (e.g., the embryonic neural tube). In contrast, cooperative mechanisms of elongation typically involve signal-based and force-based interactions. Signal-based interactions occur when the secretion of biochemicals from one tissue induces elongation in a neighbouring tissue, e.g., signalling from the ectodermal ridge induces a cell division pattern in the underlying mesenchyme to elongate the developing limb [3, 4, 5]. Force-based interactions occur when the force generated by one tissue drives elongation in neighbouring tissues, e.g., muscle-driven stretching of the *C. elegans* epidermis [6, 7, 8]. Understanding how tissues elongate autonomously or cooperatively to produce robust, elongated morphologies is a central question in developmental biology with implications for tissue engineering and regenerative medicine.

Intestinal villus morphogenesis is a model system for elongation via cooperating tissues. Intestinal villi are finger-like projections that line the surface of vertebrate guts and emerge from initially flat sheets of cells during development. Intestinal villi drastically increase the gut’s surface area, allowing nutrients to be absorbed more efficiently [9, 10]. Villus morphogenesis is not only critical for increasing the surface area of the gut but also for subsequent morphogenesis of intestinal crypts between each villus that are necessary for maintaining the digestive capabilities of the gut [11]. The morphogenesis of intestinal villi consists of two distinguishable, albeit mechanically continuous, stages: an initial folding stage and a subsequent elongation stage. During folding, intestinal tissues buckle into a stereotyped pattern of folds. During elongation, each fold transforms into its final finger-like structure. Both of these stages rely on tightly coordinated interactions between distinct tissue layers (as we soon describe), making the developing villus an ideal model for understanding how tissues cooperate to drive elongation.

Existing research has focused on the genetic pathways and cellular rearrangements involved during the initial folding stage to decipher the mechanisms driving intestinal villus morphogenesis. These pathways and rearrangements differ between avians and mammals. In mammals, folding occurs through the cooperation of an overlying epithelial tissue lining the gut surface and an underlying mesenchymal tissue. Folding begins with morphogen-induced signalling from the epithelia, causing the initially homogeneous mesenchyme to separate into an upper layer and a lower layer (Fig. 1A). The upper layer consists of densely packed, liquid-like cells, hereafter termed “L-cells,” which are marked by high levels of platelet-derived growth factor receptor alpha (PDGFRA). The lower layer consists of loosely packed, solid-like mesenchymal cells, termed “S-cells,” embedded in an extracellular matrix (ECM) termed. S-cells are marked by low levels of PDGFRA [12, 13]. The L-cells are liquid-like because they produce metalloproteinases that degrade the stiff ECM that surrounds them and express adhesion proteins that allow cells to stick together [14, 15]. Conversely, the S-cells are solid-like tissue because they do not degrade the ECM. The difference in adhesive properties between L-cells and S-cells then causes the L-cells to “dewet” from the S-cells. Dewetting is the process in which a liquid film retracts from a solid surface due to unfavourable adhesion forces between the liquid and the solid, forming isolated droplets rather than a continuous layer [16]. Dewetting causes L-cells to group into distinct, periodic clusters, pushing the epithelial layer into a pattern of folds.

**Figure 1:**
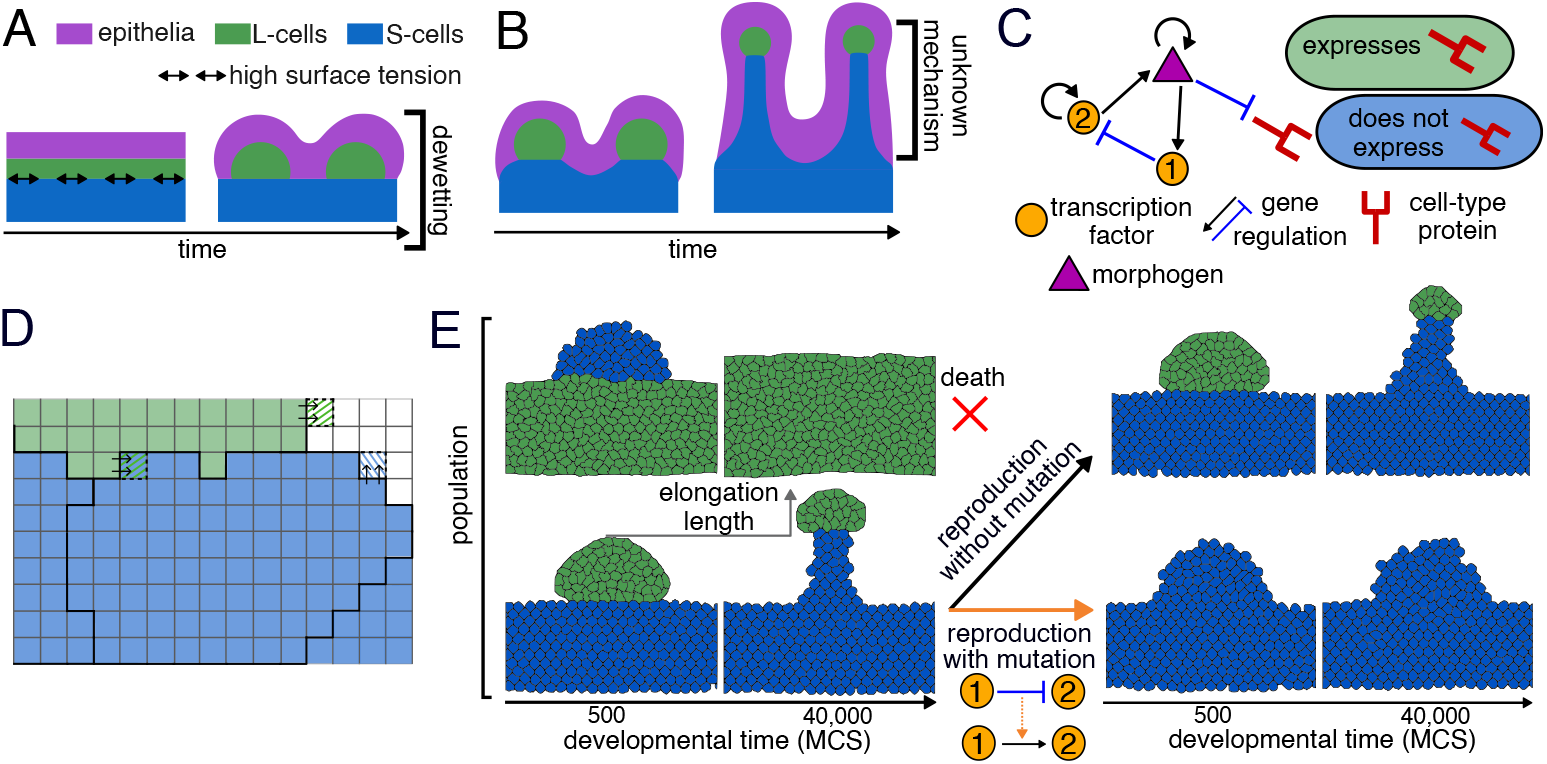
Evolutionary-developmental model of intestinal villus elongation. **A** Depiction of the onset of villus morphogenesis in mammals. Morphogenesis begins with the patterned folding of epithelial tissue (purple) due to the dewetting of liquid-like upper mesenchyme into clusters (L-cells, green) from solid-like lower mesenchyme (S-cells, blue). Dewetting occurs due to a high surface tension at the interface between L-cells and S-cells (double-sided arrows). **B** Depiction of villus elongation in all vertebrates [13]. The mesenchyme and epithelia undergo elongation to create finger-like projections, with L-cell clusters remaining at the mesenchyme’s tip throughout. **C** Example of a gene regulatory network (GRN) in our model. Each cell contains a GRN encoding three transcription factors (TFs; circles and triangle) and one protein (red stick) that determines the cell type: L-cell or S-cell. The triangle represents a membrane-permeable TF (the morphogen) that diffuses between cells. Arrows indicate the regulation of gene expression by TFs (black for activation, light blue for inhibition). **D** Close-up of the Cellular Potts Model (CPM) grid. The grid shows four neighbouring cells, with black lines depicting cell boundaries. Green colouring depicts cells in the L-cell state, blue depicts cells in the S-cell state, and white depicts the surrounding medium. Pixels with alternating stripes depict pixel copies at cell boundaries. **E** Evolutionary model consisting of a population of 60 morphologies (only two depicted for each generation). Each morphology undergoes a developmental phase on a separate CPM grid for 40,000 Monte Carlo Steps, followed by a reproduction phase in which morphologies are selected based on their elongation length (grey arrow), with the remaining morphologies dying. Reproduction can occur without mutation (black arrow) or with mutation (orange arrow), with these mutations determined probabilistically. Mutations change the topology of the GRN (dashed orange arrow). The example shows a change from inhibition of TF-2 by TF-1 to activation of TF-2 by TF-1.

During the elongation stage of villus morphogenesis, the L-cell clusters persist at the tip of each villus just underneath the epithelial cell layer (Fig. 1B) [13]. Concurrently, the epithelial cells undergo shape changes that depend on signal- and force-based interactions with the mesenchyme [17, 18]. Thus, in mammals, the mesenchyme drives folding and is likely involved in elongation as well. In avians, folding occurs through an entirely different mechanism involving tensile stress from smooth muscle contractions [19]. Mesenchymal clusters of L-cells still do appear at the beginning of the elongation stage in avians, suggesting avians and mammals might share a common elongation mechanism involving these clusters [9, 17]. However, the mechanisms of villus elongation remain unresolved.

Here, we investigate possible mechanisms of elongation by computationally modelling villus morphogenesis. We extend previous adhesion-based models of mesenchymal cells during intestinal villus folding [13] by introducing gene regulatory networks (GRNs), cell-cell signalling, and GRN-mediated transitions between L- and S-cell states to a Cellular Potts Model. To facilitate the discovery of elongation mechanisms that may not be intuitively obvious, we simulate an evolutionary algorithm that optimises for tissue elongation through mutation and selection of GRNs. We find that different evolutionary simulations converge on a common elongation mechanism involving the dewetting of L-cell clusters from S-cells, just like in the buckling phase. However, in addition to dewetting, L-cells progressively differentiate into S-cells at the interface between L-cells and S-cells. This differentiation drives the L-cell cluster to continuously dewet from the S-cells in the opposite direction of the interface, ultimately creating finger-like projections. Thus, our modelling reveals that dewetting, observed during the initial buckling of intestinal villi, can also drive elongation when coupled with differentiation.

## 2 Results

### 2.1 Evolutionary-developmental model of gut villus elongation

To investigate possible mechanisms of villus elongation, we formulated a model consisting of a population of mesenchymal cells that can switch between two states: the liquid-like “L-cell” state and the solid-like “S-cell” state. We assume that the surface tension between L-cells and S-cells is high and L-cells are more fluid-like than S-cells, in accordance with experimental observations [13]. Cells switch back and forth between states depending on the expression pattern of a “cell-state gene.” The cell is an L-cell when the cell-state gene is expressed and an S-cell when it is not. The expression pattern of the cell-state gene is determined by a gene regulatory network (GRN, described subsequently). We implement cells on a two-dimensional grid of 200 × 300 pixels, with each cell composed of a collection of neighbouring pixels. Pixels not occupied by cells represent the medium (Fig. 1D), which models the epithelial layer in between the mesenchyme and the intestinal lumen [20]. Because our focus is on mesenchymal dynamics, we treat this medium as a coarse-grained approximation of the epithelial layer rather than explicitly modelling individual epithelial cells. We use the Cellular Potts Model (CPM) formalism to simulate stochastic extensions and retractions of the cell boundaries, mimicking natural cell motion and geometry changes (Appendix 4.1) [21, 22, 23]. These extensions and retractions occur via “pixel copies,” in which the state of a random pixel on the grid replaces the state of a random neighbouring pixel. Pixel copies minimise free energy contributions from adhesion at cell boundaries and deviations from a target cell size using a probabilistic Metropolis algorithm (Appendix 4.1). Adhesion energies at cell boundaries dictate both surface tensions and cell fluidities in our model. For the results presented in the main text, we modulate the adhesion energies to ensure that the surface tension between L-cells and S-cells is a constant, high value so that L-cells are more fluid than S-cells. We test the effect of changing both the surface tension and cell fluidities in Text S1 and Fig. S1. The size of each cell, measured in pixels, is energetically constrained to a target size of 80 pixels: the more a cell’s size departs from this target size, the larger the free energy. Pixels on the grid are chosen in a random order with replacement for pixel copies. Time is measured in pixel copy attempts, with one unit of time occurring when the number of pixel copy attempts equals the total number of pixels on the grid, herein referred to as a Monte Carlo step (MCS).

To allow cells to change their state over time, we couple the CPM dynamics to the GRN, as previously done [24, 25, 26]. A GRN is a graph consisting of nodes representing proteins and edges representing activation or inhibition of gene expression (Fig. 1C). Protein concentrations within a cell are updated over time via coupled differential equations with parameters determined by the GRN (Appendix 4.2). Thus, protein concentrations are unique for each cell on the CPM grid despite all cells having the same GRN. The GRN contains four protein-encoding genes: three encoding transcription factors (TFs) and one encoding the cell-state protein. One of the three TFs diffuses between cells and into the medium, modelling a morphogen involved in intestinal villus morphogenesis, such as BMP4 [15] (Appendix 4.2). This morphogen enables cells to alter gene expression in a spatially dependent manner. We note that these GRNs do not model real villus cell GRNs; they are implemented solely to explore possible mechanisms of elongation.

Our model simplifies gut morphogenesis by focusing on a single villus, thereby avoiding the complexity of simulating the thousands of villi that form in parallel [9]. The model’s initial condition replicates early villus structure observed following mesenchymal cell aggregation: a semicircle of cells positioned on top of a flat sheet of cells. The protein encoded by the cell-state gene is initially set to a concentration of zero in all cells, meaning that all cells in both the sheet and the cluster start as S-cells. To incorporate an initial difference in gene expression between these two regions, we initialise the two non-morphogen TFs to distinct concentrations based on each cell’s initial position. One of these TFs is concentrated only in the cells starting in the flat sheet; the other TF is concentrated only in the cells starting in the semicircular cluster. Each cell begins with a size of approximately 80 pixels. For simplicity, we employ periodic boundary conditions, with the flat sheet of cells wrapping around the vertical boundary. We simulate the development of a villus for 40,000 MCS, unless otherwise specified, because this is long enough for any cell dynamics that may drive elongation to be exhausted.

To simulate evolution, we establish a population of 60 CPM grids in parallel, with each CPM grid simulating the development of one villus for the 40,000 MCS. To allow for GRN evolution, the 15 villi that undergo the largest vertical elongation are then each selected to reproduce four times to populate the next generation (Fig. 1E, Appendix 4.3). Between generations, we mutate the edges of the GRN (Fig. 1E). Each simulation runs for 10^3^ generations.

### 2.2 Repeated evolution of an elongation mechanism

To investigate possible mechanisms of villus elongation, we conducted five independent evolutionary simulations of our model. In the initial population of each simulation, each villus starts with a unique, randomly generated GRN. The results show that elongation length increased significantly over generations in all five evolutionary simulations (Fig. 2A and Fig. S4A), indicating that mechanisms driving villus elongation evolved in our model.

**Figure 2:**
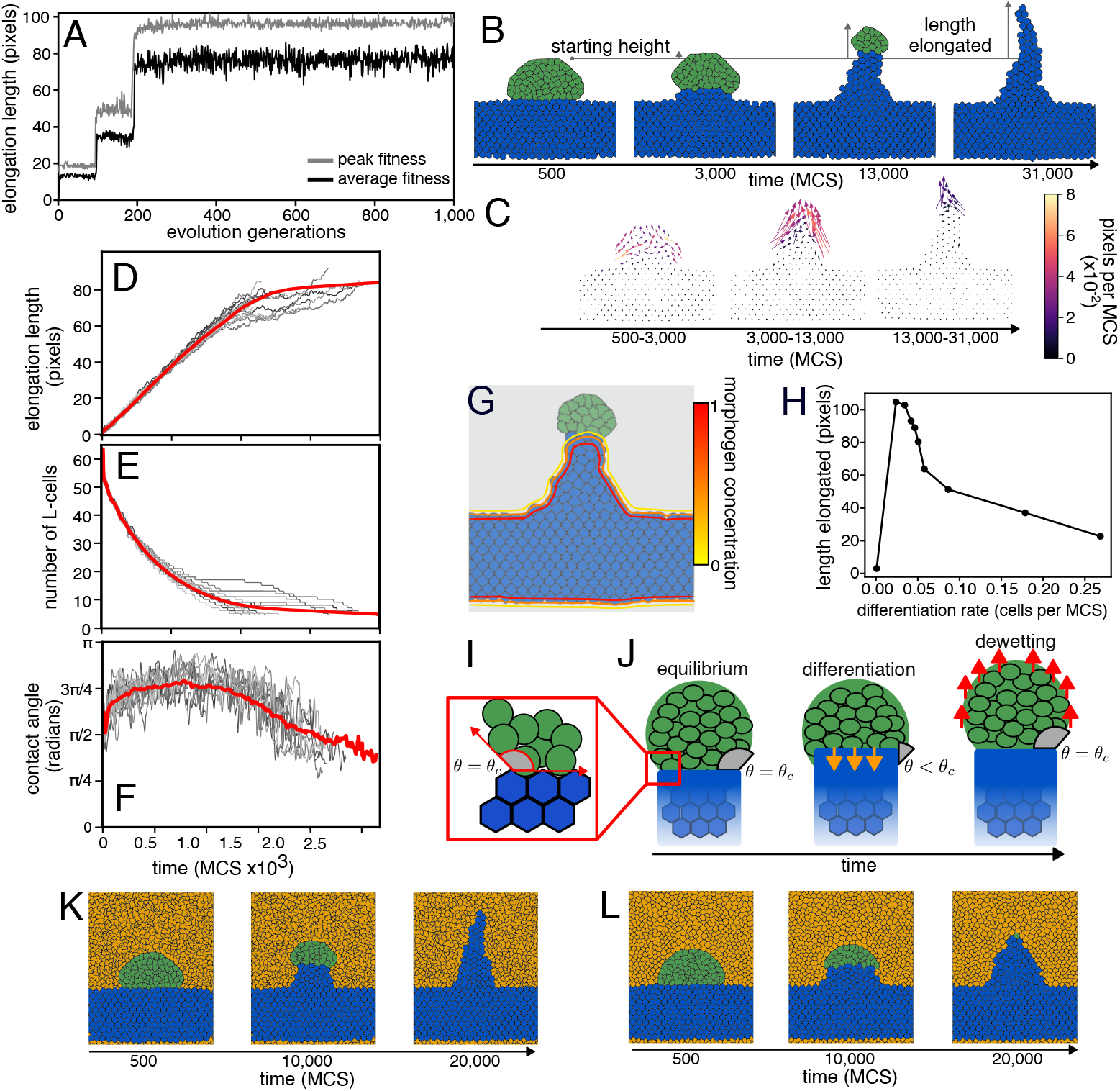
Evolution of a villus elongation mechanism. **A** Evolution of elongation lengths over a simulation. The grey and black lines represent the maximum and average elongation lengths across generations, respectively. **B** Development of an evolved morphology, with snapshots at 500, 3,000, 13,000 and 31,000 MCS. The vertical grey arrows indicate each snapshot’s elongation length from its starting height. **C** Displacements of cell centroids between each snapshot shown in (B). Arrows depict the direction of cell displacement, with their thickness scaling with speed. The colour depicts each cell’s speed in pixels per MCS. **D** Elongation length as a function of time for the evolved morphology. The red line averages 120 developmental replicates (10 are shown in grey; the rest are hidden). **E** The number of L-cells present as a function of time, using the same data as in (D). **F** Contact angles as a function of time, using the same data as in (D). **G** Snapshot of evolved gut villus morphogenesis after 15,000 MCS overlaid with a contour plot of morphogen concentration. **H** Elongation length as a function of the differentiation rate. Black dots show the differentiation rate and elongation lengths averaged over 60 developmental replicates for ten different morphogen production rates (see Appendix 4.2 and Fig. S6 for details). The differentiation rate is the number of L-cells that differentiate per MCS, measured in the first half of the total simulated time. Simulations stop when all L-cells differentiate. **I** Schematic of the contact angle (*θ*, shaded grey). Red arrows indicate tangents to the L-cell/medium and L-cell/S-cell interfaces. *θ*_*c*_ is the equilibrium contact angle determined by the balance of interfacial tensions. **J** Illustration of differentiation-induced dewetting driving elongation. The red box indicates the region around the triple point, magnified in (I). Orange arrows indicate L-cell differentiation to S-cells. Red arrows indicate L-cell motion due to dewetting. **KL** Development of an evolved morphology in the presence of an epithelial layer with (K) low viscosity, and (L) high viscosity. The epithelial cells are coloured orange.

To understand the mechanisms driving elongation, we examined the morphologies generated by the most common GRN in the final generation of each evolutionary simulation (referred to as evolved morphologies). We examined these morphologies by observing snapshots, videos and cell velocity fields (Fig. 2AB; Fig. S4B-E; Video S1). To track cell velocities, we recorded the location of cell centroids (centres of mass) for every cell in the morphology at each MCS. Interestingly, we found that all evolved morphologies displayed the following three characteristics. First, the flat cell sheet consists of S-cells. Second, at the tip of each villus is a cluster of L-cells that move in the opposite direction to the flat sheet. Third, L-cell clusters gradually disappear as they move. L-cell clusters disappear because L-cells transition to S-cells, with these S-cells remaining stationary and forming a finger-like stalk (Fig. 2AB). Since L-to-S-cell transitions appear to be irreversible, we henceforth refer to them as L-cell differentiation. Together, these three characteristics imply that elongation depends on the presence of L-cells. To confirm this, we plotted the elongation length and the number of L-cells as a function of time. We averaged these plots over 120 “developmental replicates,” which are repeat simulations using the same GRN but with unique numerical seeds (yielding different sequences of pixel copies). The results show that the tissue elongates linearly with time until the number of L-cells diminishes, after which elongation lengths plateau (Fig. 2CD shows the results for one representative evolved organism). Together, these results suggest that all evolved morphologies elongate through a similar mechanism(s) involving persistent, directional motion of L-cell clusters and L-cell differentiation into S-cells.

We hypothesised that L-cell differentiation is caused by cell-cell signalling via morphogens. To investigate this, we plotted contours of equal morphogen concentration overlaid on the morphology (Fig. 2E). The results show the morphogen concentration is high around S-cells and low everywhere else (Fig. 2E), indicating that S-cells express the morphogen whereas L-cells do not. Next, we examined regulatory interactions among genes in the GRNs of evolved organisms to see whether morphogen expression affects cell state (Fig. S4). The results show that the morphogen inhibits the expression of the cell-state gene in all five evolved GRNs. Given that cell-state gene expression causes cells to take on an L-cell state, this regulatory logic implies that L-cells differentiate only when they sense the morphogen expressed by S-cells. As a consequence, only L-cells that are close to S-cells should differentiate, which is what we observe (Video S1). Thus, S-cells induce L-cell differentiation.

To understand the relationship between L-cell differentiation and elongation, we measured elongation as a function of the L-cell differentiation rate. We focused on a single representative evolved morphology, since L-cell differentiation occurs through the same morphogen-mediated mechanism in all five. We varied the L-cell differentiation rate by changing the morphogen production rate, with higher production corresponding to faster differentiation (Appendix 4.2). The results show that elongation depends non-monotonically on the L-cell differentiation rate, with the elongation length falling if the differentiation rate is too high or too low (Fig. 2F). A non-monotonic relationship suggests that elongation is not driven solely by L-cell differentiation, but also involves additional properties.

We hypothesised that the additional property driving elongation is tissue dewetting, in which L-cells dewet collectively from S-cells, just like the buckling stage of villus morphogenesis [13]. To determine whether the L-cell clusters dewet from S-cells, we measured the contact angle, denoted *θ* (see Appendix 4.4 for a full explanation and Text S1 for validation). *θ* is the angle between the surface tangent on the L-cell/medium interface and the tangent on the L-cell/S-cell interface at the triple point where the L-cells, S-cells and medium meet (Fig. 2I). The contact angle indicates dewetting when it exceeds *π/*2. We tracked the contact angle over time across 120 developmental replicates of the evolved morphology. Since each L-cell cluster forms two distinct triple points with S-cells and the medium (one at each corner), we averaged measurements from both triple points per replicate. The results show that the contact angle rises above *π/*2 radians at the start of development and remains above this value until elongation no longer occurs (Fig. 2F). The fact that the contact angle remains above *π/*2 during villus elongation indicates that the L-cell cluster dewets from S-cells during elongation.

Our results so far lead us to hypothesise the following mechanism driving elongation. L-cell differentiation increases the interface size between L-cells and S-cells—the “SL-interface” size. Expanding the SL-interface increases energy arising from surface tension, triggering the L-cell cluster to dewet from S-cells to keep the SL-interface small. This dewetting requires L-cells to move in the direction opposite to differentiation, which elongates the morphology (Fig. 2J). Our hypothesis thus predicts that high surface tension between L-cells and S-cells is necessary for elongation, as it drives dewetting. To test this prediction, we simulated the evolved morphology with decreased surface tension between L-cells and S-cells (Text S1). The results show that decreasing surface tension yields shorter, wider morphologies (Fig. S1), confirming our prediction. In actual villus elongation, the mesenchymal cells are overlain by epithelial cells. To test whether the presence of an epithelial cell layer inhibits elongation, we replaced the medium with a large epithelial cell layer. Our results show that elongation still occurs provided epithelial cells are not too rigid (Fig. 2; see Fig. S7 for details). Together, our results indicate that the coupling of differentiation and dewetting can drive the morphogenesis of finger-like villi, hereafter termed the *dewet-differentiation mechanism*.

### 2.3 Continuum model of villus elongation

To validate our hypothesised dewet-differentiation mechanism and test how villus morphogenesis depends on biophysical determinants, including surface tension, viscosity and differentiation rates, we developed a continuum model. We shifted from the evolutionary-developmental model to a continuum model owing to several limitations of the CPM, such as constraints on tissue viscosity and non-physical behaviour at high surface tensions (see Texts S1-3 for further analysis of the CPM and its limitations). Moreover, the evolutionary-developmental model made specific assumptions about morphogens and GRNs that do not necessarily reflect those involved in real intestinal villus morphogenesis. The continuum model we developed to validate the dewet-differentiation mechanism is based on the phase-field method. The phase-field method has proven a powerful tool for describing interface-driven interactions between tissues [27, 28, 29, 13, 30]. In the continuum model, we treat scalar fields as representing distinct tissues that average the behaviour of cell populations. Our continuum model avoids the limitations of the CPM and makes no specific assumptions about the morphogens and GRNs involved in intestinal villus morphogenesis.

The continuum model describes the three tissue types on a 4 *×* 6 domain with periodic boundary conditions: L-tissue, S-tissue and E-tissue, representing L-cells, S-cells and epithelial cells, respectively. Each tissue is represented by a concentration between one and zero: *c*_*L*_, *c*_*S*_ and *c*_*E*_. These concentrations are constrained to sum to one at every point on the domain, i.e., *c*_*L*_ + *c*_*S*_ + *c*_*E*_ = 1. Each tissue consists of bulk and interface regions. For a given tissue, a concentration equal to 1 signifies the tissue bulk, while a concentration equal to 0 signifies that the tissue is not present. The transition between a concentration of 1 and 0 occurs smoothly across a narrow tissue interface region (Fig. S8AB). The width of the interface region is constant across all possible tissue interfaces and simulations (Appendix 5).

We formulate an initial condition of our continuum model that resembles a single intestinal villus following L-cell aggregation into clusters. Specifically, we initialise our model to a flat layer of S-tissue with a semicircle of L-tissue on top (See Text S1 for equations). To test the robustness of our results, we also simulate our model from an initial condition that resembles a single intestinal villus prior to L-cell aggregation (Fig. S9, Text S1).

We model L-tissue, S-tissue and E-tissue as viscous fluids. To reflect findings that S-tissue behaves as a solid while L-tissue behaves as a liquid on developmental timescales [13], we initially set the relative viscosity of S-tissue to L-tissue, termed *η*, equal to 100. High S-tissue viscosity ensures that it deforms slowly, approximating solid-like behaviour on the timescale of elongation.

We assume that tissues exist in a dissipative environment, with tissues changing their shape and location over time to minimise free energy arising from surface tension at tissue interfaces (in arbitrary time units *t*). Surface tensions are determined by parameters *γ*_*SL*_, *γ*_*SE*_ and *γ*_*LE*_, which denote the surface tension at the SL-interface, SE-interface and LE-interface, respectively (Appendix 5). Surface tensions determine the equilibrium shapes of tissues, i.e., whether the L-tissue dewets from S-tissue or wets over S-tissue (Fig. S8). The equilibrium tissue shapes produced by our model agree with predictions using classical wetting theory (see Fig. S10 for examples).

We introduce differentiation of L-tissue into S-tissue into our model with no specific biological assumptions about the cause of differentiation, except that it occurs at the interface between the two tissues. Therefore, this differentiation can also represent the physical process of L-cells detaching from the L-tissue cluster and sinking into S-tissue extracellular matrix, a phenomenon thought to occur during intestinal villus morphogenesis [18]. The differentiation rate per contact length at the SL-interface is determined by a parameter *D* (Appendix 5).

Simulations of our model last until one of the following conditions is reached: (i) the volume of L-tissue drops to zero because it all differentiates into S-tissue, or (ii) L-tissue separates from S-tissue since the mesenchyme must remain intact to generate intestinal villi during real morphogenesis [18]. To quantify elongation in our model, we measure the difference between the maximum vertical extent of the S-tissue (defined as the highest *y*-coordinate where *c*_*S*_ *>* 0.5) at the simulation endpoint and its maximum vertical extent at *t* = 0, termed the “elongation length.”

### 2.4 Balance between dewetting and differentiation determines villus morphologies

We asked how tissue surface tensions, and thus dewetting, shape villus morphologies in our continuum model. To answer this, we simulated the model over a parameter sweep of *γ*_*SL*_ and *γ*_*LE*_, while setting *D* = 1 and *γ*_*SE*_ = 0.06. We chose this value of *D* because it corresponds to differentiation- and surface tension-driven dynamics occurring at similar timescales across our parameter sweep. We chose this value of *γ*_*SE*_ = 0.06 so that the parameter sweep spans all classical “spreading” regimes [16], where energy arising from surface tension is minimised by one tissue forming a thin, intervening layer between the other two. Spreading is predicted to occur when one surface tension exceeds the sum of the other two, e.g., E-tissue spreads between S- and L-tissue when *S*_*E*_ = −*γ*_*SL*_ *γ*_*SE*_ −*γ*_*LE*_ *>* 0, where *S*_*E*_ is the “spreading coefficient” of E-tissue. Our parameter sweep also spans the “non-spreading” regime, which lies in between the three spreading regimes. Our results show that when *γ*_*SL*_ *< γ*_*LE*_ (in or close to the S-tissue spreading regime), elongation lengths are small, with villus morphologies having thick, triangular stalks (Fig. 3AE). When *γ*_*SL*_ ≈ *γ*_*LE*_ (non-spreading regime), the S-tissue undergoes extensive elongation, giving rise to thin, elongated stalks that resemble intestinal villi (Fig. 3BG, Video S2). When *γ*_*SL*_ *> γ*_*LE*_ (close to or in the E-tissue spreading regime), there is little or no elongation.

**Figure 3:**
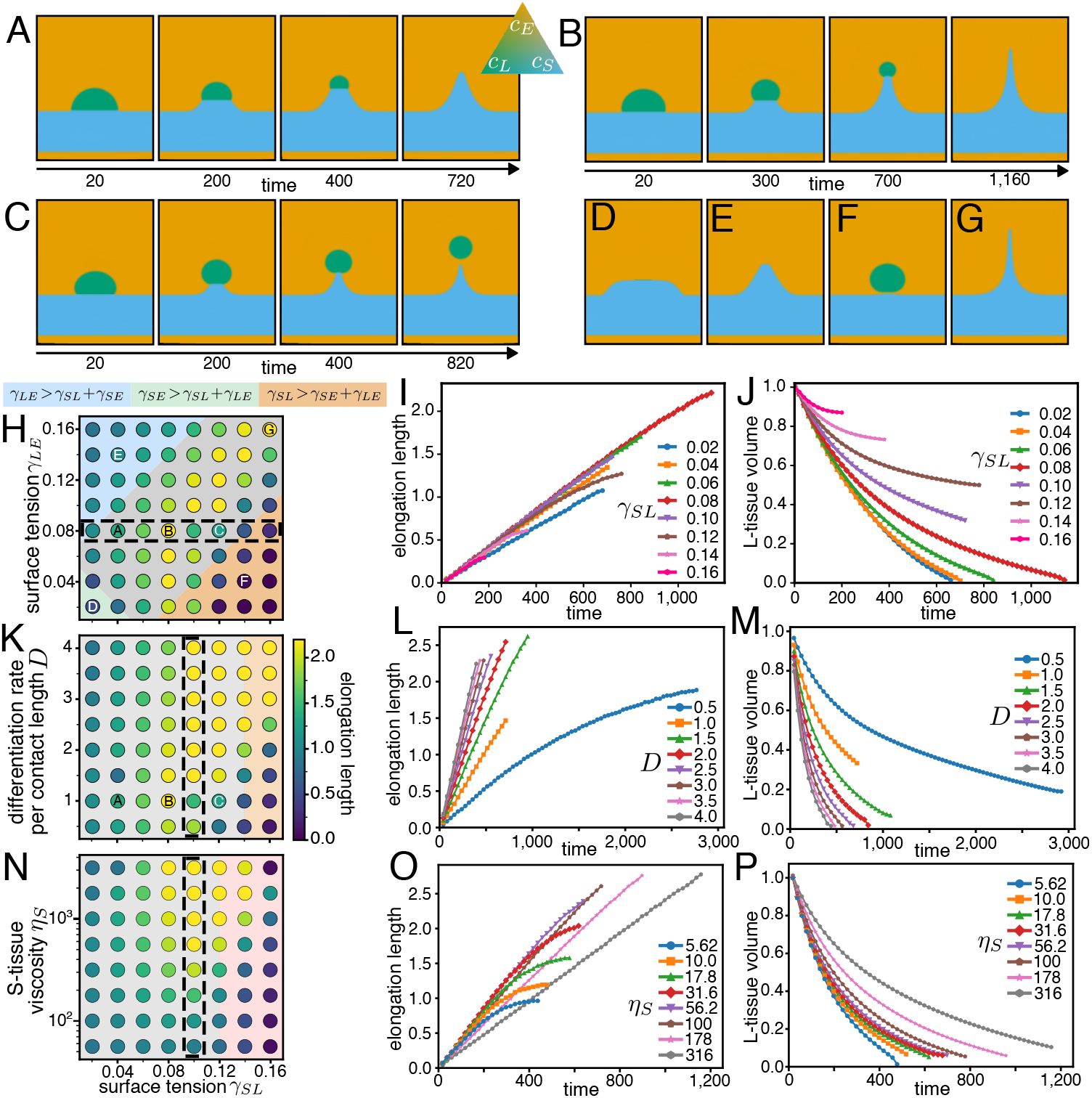
Analysis of villus elongation in the continuum model. **A**,**B**,**C** Snapshots of three elongating villi at consecutive time-points in a simulation, with time-points shown below the snapshot. Snapshots are coloured by the concentration of *c*_*S*_, *c*_*L*_, and *c*_*E*_ following the colour mapping of the ternary plot. The colour at each vertex of the ternary plot represents a pure tissue, i.e., *c*_*S*_ = 1, *c*_*L*_ = 1, or *c*_*E*_ = 1. The parameters used are *γ*_*SE*_ = 0.06, *γ*_*LE*_ = 0.08, *D* = 1 and *γ*_*SL*_ = 0.04 in (A); *γ*_*SL*_ = 0.08 in (B); and *γ*_*SL*_ = 0.12 in (C). **D-G** Snapshots of simulation endpoints for four simulations in the (D) L-tissue spreading regime, (E) S-tissue spreading regime, (F) E-tissue spreading regime, and (G) non-spreading regime. The snapshots are taken when: (D): *t* = 360, (E): *t* = 580, (F): *t* = 300 and (G): *t* = 1, 500. See the caption for (H) for parameters. **H**,**K**,**N** Phase diagrams showing simulation-end elongation length (circle colour, see heatmap) as a function of *γ*_*SL*_ and (H) *γ*_*LE*_, (K) *D*, and (N) *η*. The background colour of the phase diagram indicates the spreading regime, as indicated on top (blue is where S-tissue spreads (**S**_*S*_ *>* 0), green is where L-tissue spreads (**S**_*L*_ *>* 0), and orange is where E-tissue spreads (**S**_*E*_ *>* 0)). Grey is the non-spreading regime (**S**_*i*_ *<* 0 for *i* = *S, L, E*). **B**,**C**,**E**,**F**,**H**,**I** Temporal dynamics of elongation length and L-tissue volume for parameters corresponding to the dashed boxes in their respective phase diagrams to the left. Panels (B, C) vary *γ*_*SL*_; (E, F) vary *D*; and (H, I) vary *η*. Fixed parameters: *γ*_*SE*_ = 0.06 throughout; *D* = 1 in (H-J); *η* = 100 in (H-M); *γ*_*LE*_ = 0.08 in (K–P); and *D* = 2 in (N–P).

The lack of elongation is caused by L-tissue separating from S-tissue, resulting in thin but short villi near the boundary of the E-tissue spreading regime (Fig. 3C) and no villi at all inside the E-tissue spreading regime (Fig. 3F). When both *γ*_*SE*_ and *γ*_*SL*_ are small (L-tissue spreading regime), the L-tissue flattens out over the S-tissue surface before differentiating, resulting in a wide bulge of S-tissue (Fig. 3D). Together, our results indicate that surface tensions can substantially alter the width and height of villus morphologies, and that finger-like villi might arise only when there is a balance of surface tensions at the different tissue interfaces.

We next asked why villi with different surface tensions differ in length. The final length of a villus depends on its elongation speed and how long it takes for all of the L-tissue to differentiate (or separate from S-tissue). Thus, we tracked the elongation length and the L-tissue volume over time across the parameter sweep shown in Fig. 3H. The results show villi elongate at approximately the same speed regardless of *γ*_*SL*_ or *γ*_*LE*_ (Figs. 3I and S11B). In contrast, the L-tissue volume decreases more rapidly as *γ*_*SL*_ increases (Fig. 3J) or *γ*_*SE*_ decreases (Fig. S11B). In our continuum model, the rate of L-tissue loss via differentiation scales with the SL-interface width and *D*. Since *D* is constant in these simulations, our results suggest that elongation is governed by the SL-interface width, which is driven to an energetically preferred width by surface tensions. For instance, in or near the S-tissue or L-tissue spreading regimes, a long SL-interface width is favoured, which results in rapid L-tissue differentiation and thus shorter elongation lengths (Fig. 3H). Conversely, in the E-tissue spreading regime (e.g., high *γ*_*SL*_), surface tensions drive a rapid dewetting that shrinks the interface faster than L-tissue differentiates, eventually causing L-tissue to separate from S-tissue. Based on these results, we predict that elongation lengths are maximised when the SL-interface is narrow enough to prevent rapid L-tissue volume loss due to differentiation, yet wide enough to prevent separation of L-tissue from S-tissue due to surface tension, i.e., differentiation and surface tension are balanced.

To test our prediction that a balance of differentiation and surface tension maximises elongation, we simulated the model across a parameter sweep of *D* and *γ*_*SL*_. The results show that high elongation lengths are concentrated along a diagonal trajectory in the parameter space (Fig. 3K). Interestingly, this diagonal band even enters the E-tissue spreading regime, where L-tissue and S-tissue are predicted to separate. To understand why this separation does not occur, we observed snapshots of morphologies in this E-tissue spreading regime. The snapshots show that fast differentiation prevents the dewetting-induced separation of L-tissue and S-tissue, resulting in finger-like morphologies (Fig. S12). Thus, we predict that finger-like villi can form outside of the non-spreading regime if *D* is high enough to suppress surface tension-induced separation of S-tissue and L-tissue. If *D* is too high relative to *γ*_*SL*_, however, final elongation lengths decrease even though elongation speeds increase (Fig. 3LM), with morphologies becoming more triangular than finger-like (Fig. S12). This result indicates that there is a point at which L-tissue differentiates faster than it can dewet, as observed in the evolutionary-developmental model (Fig. 2H). If *D* is very low, S-tissue and L-tissue can separate even in the non-spreading regime (e.g., Fig. 3C). In this case, separation occurs because stalk widths fall below the width of the tissue interface region, which is determined by a parameter in our continuum model (Appendix 5). Since the tissue interface width is an artefact of the phase-field model, the separation observed in the non-spreading regime is not biologically realistic unless the interface width is smaller than a cell diameter. Together, these results demonstrate that a balance between differentiation and surface tension at the SL-interface facilitates villus elongation.

We next asked how tissue viscosity affects villus morphology in our continuum model. To answer this, we performed a parameter sweep over *γ*_*SL*_ and the relative viscosity of S-tissue to L-tissue *η* (set to 100 in previous simulations). The results show that *η* must be sufficiently high for elongated, finger-like morphologies (Fig. 3N). When *η* ∼ 10^0^–10^1^, elongation lengths are small for all *γ*_*SL*_ because the S-tissue surface resists being stretched into a finger-like morphology that increases energy from surface tension. As *η* increases, elongation lengths also increase until *η*≈50, after which they plateau around a peak length (Fig. 3O). However, the speed of elongation declines when *η >* 100 because SL-interface widths become smaller (Fig. S13). These results show that S-tissue viscosity needs to be significantly higher than L-tissue viscosity for the dewet-differentiation mechanism to facilitate elongation.

## 3 Discussion

Sub-epithelial mesenchymal cells are thought to play a crucial role in intestinal villus elongation [15, 18], but the specific mechanism by which these cells drive elongation remains poorly understood. In this study, we explored possible mechanisms of villus elongation using an evolutionary-developmental model that simulates a population of mesenchymal cells. By evolving the morphologies of mesenchymal cells under selection for elongation, we observed the repeated emergence of a *dewet-differentiation mechanism*. This mechanism depends on two interacting mesenchymal cell populations to drive elongation: liquid-like L-cells, which form clusters at nascent villi tips, and solid-like S-cells, which reside below the cluster. This mechanism builds on recent work showing that dewetting drives the initial clustering of L-cells into periodic folds along the surface of the gut, with these folds demarcating the sites of each villus [13]. Our results show that by incorporating differentiation of L-cells to S-cells into this system, elongation can occur. This elongation is a consequence of dewetting, which reduces the interface width between L-cells and S-cells, while simultaneously L-cells at this interface differentiate into S-cells, thereby expanding the interface width (Fig. 2J). To maintain minimal surface energy, the remaining L-cells continuously dewet in the direction opposite to the S-cells, elongating the morphology. Thus, our modelling, which we theoretically validated in a continuum phase-field model, reveals that the continual induction of dewetting by differentiation can drive the transition from initial villus folding to subsequent villus elongation.

An advantage of the dewet-differentiation mechanism is its simplicity, since both dewetting and differentiation are fundamental processes of multicellular development. While direct experimental evidence confirming that dewetting and differentiation occur during villus elongation is currently lacking, existing literature provides indirect support. For instance, because dewetting occurs during the initial folding stage of villus morphogenesis [13], it is highly plausible that it continues during elongation. Similarly, indirect evidence supports L-cell-to-S-cell differentiation during elongation. As the villus elongates, cells within the L-cell mesenchymal cluster progressively disperse [9, 18]—a spatiotemporal pattern that closely mirrors differentiation in our model. Furthermore, preventing L-cell dispersal by artificially increasing cohesion within the L-cell cluster has been shown to impede villus elongation [18]. Thus, to validate the dewet-differentiation mechanism, we propose that future experimental studies test whether L-cells differentiate into S-cells during this dispersal and whether L-cells dewet from S-cells during elongation.

There are two distinct modes by which differentiation could occur in real intestinal villi. The first is biochemical differentiation, in which cell-cell signalling (driven by morphogens such as Hedgehog, Wnt, or PDGF) triggers a change in L-cell gene expression, leading them to transition into S-cells (like our Cellular Potts Model). Biochemical differentiation could be tested by pharmacologically or genetically perturbing potential morphogen pathways in intestinal explant models and observing whether this causes the L-cell cluster to transition into ECM-producing S-cells [31]. The second potential mode is mechanical ingression, wherein L-cells delaminate from their cluster and embed into the underlying ECM. This mode is consistent with our continuum model. Testing this would require live-cell imaging on intestinal explants to capture the spatial trajectories of individual L-cells [32]. Ultimately, regardless of whether differentiation is biochemical or mechanical, transcriptional profiling or genetic lineage tracing should reveal a change in gene expression from L-cell to S-cell states if the dewet-differentiation mechanism drives elongation. Specifically, we would expect to see the progressive downregulation of L-cell-associated adhesion molecules (integrins and cadherins) and ECM-degrading enzymes if L-cells differentiate into S-cells.

Some experimental observations suggest that additional mechanisms could be involved in intestinal villus elongation. For instance, the secretion of signals by L-cells during villus elongation affects the proliferation and shape of the overlying epithelial cells [15, 17], suggesting that epithelial tissue might cooperate with mesenchyme to sculpt intestinal villi morphology. Future modelling could incorporate epithelial cell behaviours, such as division, contraction, and basement membranes, to determine how epithelial behaviours interact with the dewet-differentiation mechanism to facilitate villus elongation. This model could be used to explain variations in villus morphologies observed along the gut axis [33].

The dewet-differentiation mechanism we describe here could facilitate tissue elongation in other structures, as it requires only three properties widespread in animal development: a liquid-like tissue adjacent to a solid-like one, high surface tension between these tissues, and cell differentiation from the liquid-like to the solid-like tissue. Moreover, the mechanism is both scalable and robust, with elongation speeds and morphologies changing predictably with surface tensions and differentiation rates (Fig. 3). Possible elongating structures in which the dewet-differentiation mechanism could play a role include those involving mesenchymal clusters. Mesenchymal clusters, such as those made up of L-cells in intestinal villi, satisfy the above requirements because they tend to be liquid-like, found adjacent to solid-like tissues, and often differentiate into these solid-like tissues [34, 35]. Beyond intestinal villi, mesenchymal clusters are involved in kidney nephron elongation and limb bud elongation [36, 37]. Future investigations could test whether dewet-differentiation contributes to morphogenesis in these systems.

In conclusion, we propose that the dewet-differentiation mechanism, identified through evolutionary simulation and validated via continuum modelling, is a plausible mechanism of intestinal villus elongation. This novel mechanism hinges on the interplay between tissue-scale dewetting and mesenchymal cell differentiation, is consistent with existing experiments on villus morphogenesis, and offers testable predictions for future experiments. Investigating interactions between biochemical processes, such as cell differentiation, and biophysical properties, such as dewetting, is fertile ground for developing a better understanding of the morphogenic mechanisms that underlie developmental processes, such as tissue elongation.

## Acknowledgements

The authors would like to thank Kota Mitsumoto for fruitful discussion.

## 4 Appendix A: Hybrid Cellular Potts Model

### 4.1 Cellular Potts Model

We introduce a hybrid Cellular Potts Model (CPM) by adapting an implementation of the Tissue Simulation Toolkit [38, 39]. The CPM simulates cells on a two-dimensional grid, with each cell composed of a collection of neighbouring pixels. The pixels not occupied by cells represent the medium. Cell motion occurs via stochastic “pixel copies.” Pixel copies occur by (i) selecting a random pixel on the grid, (ii) selecting a random pixel within the Moore neighbourhood of the first pixel, and (iii) replacing the state of the first pixel with the second pixel depending on the change to the system energy. The unit of time is the number of pixel copies equal to the total number of pixels on the grid, referred to as a Monte Carlo step (MCS), and is denoted by *t* in mathematical equations. Within each MCS, pixels are selected for copies with replacement (i.e., the same pixel can be selected more than once).

Pixel copies succeed if Δ*H <* 0, where Δ*H* is the change in the system energy if the pixel copy is successful. If Δ*H >* 0, the pixel copy succeeds with probability exp(−Δ*H/*𝒯), where 𝒯is a temperature-like parameter that we arbitrarily chose as 𝒯= 1 for all simulations. The system energy *H* is determined by free energy contributions from both adhesion at cell interfaces and a constraint on cell sizes, as follows:

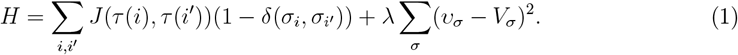

Here, *i* is an index of every pixel in the CPM, *i*′ are pixels within the Moore neighbourhood of each *i, τ*(*i*) is a function that returns the tissue type of the cell occupying pixel *i*, and *σ* is an index of every cell in the CPM, except for *σ* = 0, which represents the medium. The function *J*(*τ*(*i*), *τ*(*i*′)) takes the tissue types of cells at pixels *i* and *i*′ as arguments and outputs the adhesion energy arising between them, denoted 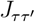 for simplicity. The function *δ*(*σ*_*i*_, *σ*_*i*_′) is the Kronecker delta, which evaluates to one when pixel *i* and *i*′ belong to the same cell *σ*, and to zero otherwise. Thus, the energy contributions from adhesion arise only at the interfaces between cells or between cells and the medium. The current size of each cell in pixels, *v*_*σ*_, is constrained to a size *V*_*σ*_ by parameter *λ* (*λ* = 1*/*6 for all simulations). We set *V*_*σ*_ = 80 for all simulations.

The adhesion energy values, 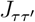, are determined by the kinds of interacting entities (*τ* and *τ* ′): L-cells, S-cells, or the medium. The interactions between these entities lead to five adhesion energies: *J*_*LL*_ (between L-cells), *J*_*SS*_ (between S-cells), *J*_*SL*_ (S-cells to L-cells), *J*_*LM*_ (L-cells to medium) and *J*_*SM*_ (S-cells to medium). Collectively, these adhesion energies determine the surface tensions at interfaces (see Text S1 for full details). The homotypic adhesion energies (*J*_*LL*_ and *J*_*SS*_) govern tissue viscosities, e.g., low *J*_*LL*_ corresponds to low viscosity in a tissue consisting of L-cells [21]. We selected adhesion energies to model low viscosity among L-cells, high viscosity among S-cells, and high surface tension at the interface between L-cells and S-cells. Specifically, we set *J*_*LL*_ = 0.66, *J*_*SS*_ = 5.17, *J*_*SL*_ = 5.25 and *J*_*LM*_ = *J*_*SM*_ = 2.33. We test the robustness of these parameters in Text S1.

### 4.2 Gene regulatory network and morphogens

In each cell *σ*, there are *N* = 5 genes, indexed by *p* ∈ [1, 5], with each gene encoding a protein whose intracellular concentration is denoted by 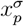. Except for *p* = 5, which is the morphogen (described later), the following equation determines the change in 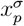over time *t*:

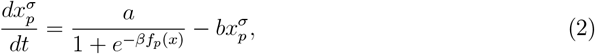

with one unit of time *t* equal to one MCS. The first term on the right-hand side represents the increase in 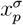 due to gene expression and is a sigmoidal function that depends on transcription factor (TF) regulation, with a maximum production rate *a* and a large *β* (= 20). The second term represents protein decay with a rate denoted by *b*. We set *a* = *b* for all *p*, so 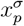 equilibrates at one if the gene is constantly expressed or zero if it is constantly not expressed. The values of *a* and *b* are small (specifically, 6.25 ×10^*−*3^) because we assume that the timescale of gene expression is slower than the timescale of CPM dynamics. Numerical integration of Equation 2 occurs with Δ*t*_1_ = 40 via the Euler method (we use a subscript because Equation 4, described subsequently, is numerically integrated at a different interval). We chose this value of Δ*t*_1_ rather than a smaller one to improve computational speed. The value of Δ*t*_1_ can be large because the rate parameters *a* and *b* are very small. The initial concentrations for transcription factors are either one or zero, depending on whether the cell is part of the flat sheet or the semicircular bulge at the beginning of development. For cells in the flat sheet, 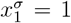 and 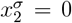. For cells in the semicircular bulge, 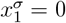 and 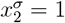. The concentration of the cell-state protein, 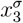, starts at a concentration of zero for all *σ*. Initial concentrations for 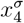 and 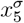 are zero (described in detail later).

Function *f*_*p*_(*x*) in Equation 2 sums the regulatory effects of the *n* = 3 transcription factors (including the morphogen) as follows:

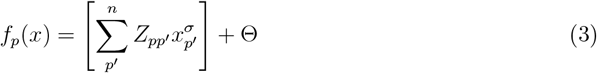

where 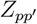 is the regulatory effect of TF encoded by gene *p*′ on the expression of gene 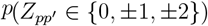. The TF encoded by *p*′ activates the expression of *p* if 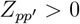, inhibits if 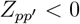 and has no effect if 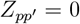. The parameter Θ sets the base level of gene expression. We set Θ = −0.3 for all simulations, so protein concentrations equilibrate at 0 when not regulated by any TFs.

To model cell-cell signalling, one of the three TFs is a morphogen, indexed by *p* = 5. The morphogen diffuses between cells and into the surrounding medium. The concentration of the morphogen on pixel *i* on the grid, 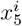, is determined by the following coupled ODE:

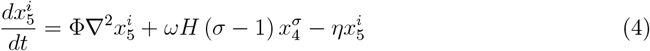

where Φ is a diffusion constant, *ω* is a production rate, *η* is a decay rate, and *H*(*σ* −1) is the Heaviside step function that evaluates to one if *σ* ≥ 1 at pixel *i* (the pixel is occupied by a cell) and zero if *σ* = 0 (the pixel is medium). The concentration 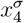 represents an intracellular signalling (IS) protein that activates the expression of the morphogen. The IS protein does not regulate other genes, whereas the morphogen does. However, the IS protein is regulated by other genes, whereas the morphogen is not. Thus, the IS protein and the morphogen can be thought of as a single node in the GRN, as described in the main text. 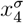 is determined by Equation 2 and starts with a concentration of zero in all cells. The operator 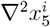 is the Laplacian acting on 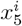, which, in this context, is the difference between 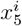 and the average 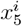 in its von Neumann neighbourhood (i.e., four nearest-neighbouring pixels) divided by a space step (*dx*)^2^, with *dx* = 1*/*250. Numerical integration for Equation 4 occurs with Δ*t*_2_ = 1. Equation is subject to the initial condition 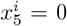 for all *i* pixels. The pixels at the boundary of the grid are subject to 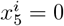 for all *t*. The constants Φ = 8 *×* 10^*−*7^, *ω* = 3 *×* 10^*−*3^ and *η* = 2 *×* 10^*−*3^were used for all evolutionary simulations, but *ω* is tuned to change the rate of differentiation in subsequent simulations. We chose these values for two reasons. The first is so that the maximum concentration is approximately one and thus similar to the concentrations of TFs per Equation 2. The second is so that the characteristic diffusion length, 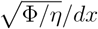, is similar to that of a paracrine morphogen, such as Wnts, which localises signalling to only nearby cells [40]. This characteristic diffusion length is approximately five pixels, which is equivalent to a cell’s radius. To obtain 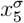 in each cell *σ* from the grid of 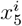 concentrations, we average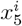 over all *i* occupied by cell *σ*.

The concentration of the cell-state protein, 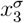, determines whether cell *σ* is an S-cell or L-cell. Since 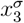 rapidly converges to zero or one, depending on the concentrations of TFs and the specific GRN (see Eq. 2, we binarise the cell state using a threshold of 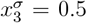. Specifically, cell *σ* is classed as an L-cell when 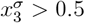 and as an S-cell when 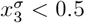.

### 4.3 Evolution

To find GRNs that cause morphologies to elongate, we ran evolutionary simulations of our CPM and evolved the regulatory effects of the GRN. We established an initial population of 60 morphologies in each evolutionary simulation, with each morphology assigned a different GRN and developing on a separate CPM grid. Each GRN is specified by 12 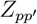 values representing the regulatory effects of all transcription factors, as described in Equation 3 (three transcription factors regulate four genes, including themselves). The 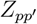 values of the GRNs assigned to the initial population are randomly generated according to the following probabilities: 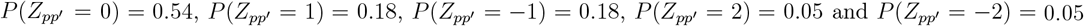. The 15 morphologies with the greatest elongation length reproduce four times to populate the next generation. Upon reproduction, there is a 50% chance that one (and only one) of the 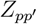 values in the GRN mutates. The specific 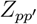 that mutates is chosen at random with equal probability. The mutation alters the value of 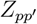 independently of its current value, according to the same probability distribution used to generate the 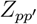 values in the initial population. The initial configuration of cells, the number of genes, *J*(*τ*(*i*), *τ*(*i*′)) values, and the types of genes remain unchanged throughout evolution. Although *J*(*τ*(*i*), *τ*(*i*′)) values do not mutate, we explore their effect on morphogenesis in Figs. S1 and S5.

### 4.4 Contact angle Measurement

We measured the contact angle over time *t* in the CPM (shown in Fig. 2F) as follows. We located the centroids of the L-cells at the two triple points where L-cells, S-cells, and the medium meet. We denote these centroids as *p*_*Q*_(*t*) and *p*_*Q*′_ (*t*), where *Q* and *Q*′ denote the L-cells at the first and second triple points, respectively. Typically, the first triple point is on the left and the second is on the right. Next, we locate the coordinates of the centroids of the L-cells that neighbour *Q* and *Q*′ and also interface with the medium. We denote these centroids as *p*_*q*_(*t*) and *p*_*q*′_ (*t*), where *q* and *q*′ denote the L-cells neighbouring the first and second triple point, respectively. We then construct vectors between centroids *Q* to *q* and *Q*′ to *q*′: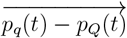for the first triple point and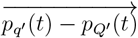 for the second. We approximate the contact angle, *θ*(*t*), as:

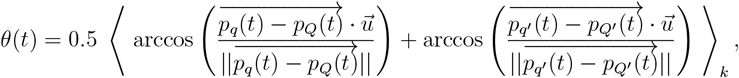

where 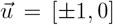 is a reference unit vector parallel to the *x* -axis whose sign is positive for the triple point on the left side and negative for the triple point on the right side. The *x* -axis is chosen as the direction for the reference vector because it runs parallel to the SL interface. The brackets ⟨…⟩_*k*_ denote an average over replicate simulations indexed by *k*. We add the prefactor 0.5 because we are summing two contact angles on either side of the morphology in each developmental replicate.

The approximations we have made to measure the contact angle are valid under two conditions: (i) that the distance between the centres of adjacent L-cells is small relative to the curvature of the interface between L-tissue and the medium, and (ii) the interface between L-cells and the medium remains parallel to the *x* -axis. The first condition results in the contact angle measurements being slightly smaller than they should be when the number of L-cells is small, which is why the contact angle decreases in Fig. 2H. In Figs. 2H and S1F, contact angle measurements stop for each developmental replicate when only a single L-cell remains in contact with S-cells, as our measurement method cannot distinguish between the two triple points when there is only a single cell remaining.

## 5 Appendix B: Continuum Model

Our phase-field model introduces *N* = 3 scalar fields, denoted generically by *c*_*i*_(**x**, *t*) (dependent on spatial coordinate **x** and time *t*), abbreviated to *c*_*i*_, with each field representing a different tissue type (*c*_*s*_ for S-tissue, *c*_*L*_ for L-tissue, and *c*_*E*_ for E-tissue). The value of each field varies continuously between *c*_*i*_ = 0 (complete absence of tissue *i*) and *c*_*i*_ = 1 (complete presence of tissue *i*). The energy generated at tissue interfaces contributes to the total free energy, *F*, that resembles the model introduced by Nestler et al (2008) [41]. Specifically, *F* is given by the following functional:

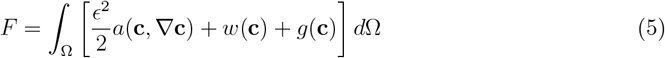

where Ω is the spatial domain, **c** = {*c*_*S*_, *c*_*L*_, *c*_*E*_}, *ϵ* is a small positive parameter that determines the thickness of the interface. The three terms inside the square brackets are the gradient energy density, the bulk potential, and the volume-constraint contributions to the free energy, respectively. While this study is limited to three tissue types, the notation used in the subsequent description of the free energy terms is intentionally general to accommodate a larger number of tissues. The model is constrained by

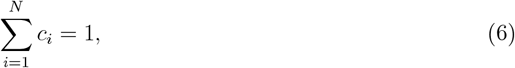

which is satisfied by setting 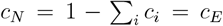. The gradient energy density, *a*(**c**, ∇**c**), establishes the surface tension between tissues by penalising sharp interfaces. It is defined as:

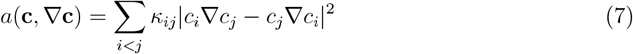

where *κ*_*ij*_ is a parameter that contributes to the surface tension, denoted *γ*_*ij*_, at the interface between tissues *i* and *j*. Specifically, *γ*_*ij*_ = *ϵκ*_*ij*_*/*3 (see Text S1 for derivation). The bulk potential, *w*(**c**), is a multi-well function that drives the system towards distinct tissue phases by energetically penalising mixtures of phases at the same location in space. It is defined as:

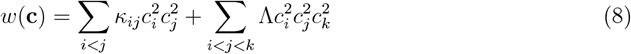

where the first term on the RHS is the double-well potential for each pair of phases and the second term on the RHS is a sextic term that suppresses non-physical third phase contributions along two phase interfaces, as used in previous multi-phase-field models [42, 41]. The exact value of the parameter Λassociated with the sextic term does not affect model dynamics as long as it is sufficiently high to suppress these third phase contributions. We set Λ= 2, 000 for all simulations. The volume constraint, *g*(**c**), is defined as:

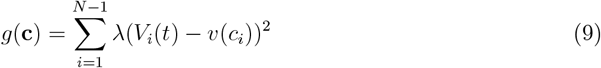

where *V*_*i*_(*t*) is the target volume of tissue *i* and *v*(*c*_*i*_) is the current volume of tissue *c*_*i*_. The values of *V*_*i*_(*t*) depend on the initial conditions of *c*_*S*_ and *c*_*L*_ and subsequent cell differentiation (explained later). We calculate *v*(*c*_*i*_) by:

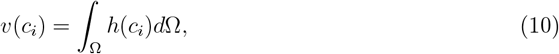

where 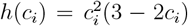. This function *h*(*c*_*i*_) is used because it more accurately calculates the geometric volume of the phase instead of an integral of *c*_*i*_ over Ω. We assume that tissues behave as over-damped fluids undergoing relaxational dynamics (i.e., model A dynamics [43], leading to the following time evolution Equations:

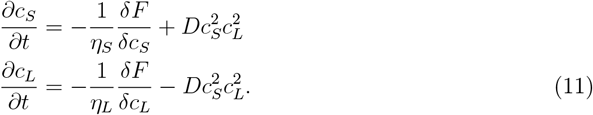

where *η*_*S*_ and *η*_*L*_ are parameters that determine the viscosity of S-tissue and L-tissue, respectively. We let *η*_*L*_ = 1 in all simulations and refer to the ratio *η*_*S*_*/η*_*L*_ as *η* in the main text for simplicity. The final term on the RHS is the differentiation term, where *D* is a parameter governing the differentiation rate from *c*_*L*_ to *c*_*S*_ at any interface between these two tissues. We chose a squared form for the differentiation term to avoid diffuse creation of S-tissue inside bulk L-tissue. The target volume of S-tissue, *V*_*S*_, increases over time while the target volume of L-tissue, termed *V*_*L*_, decreases over time due to this differentiation, as follows:

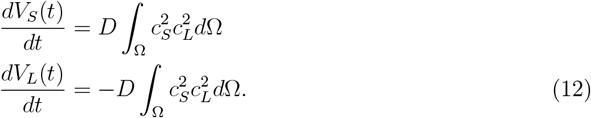

We set *D* = 0 for all simulations until *t* = 10 to ensure tissue interfaces are fully relaxed before differentiation begins.

### Numerical implementation

We implemented our model using FreeFEM++ software [44]. We solve our model on a rectangular domain spanning 0≤ *x*≤ 4 and 0≤ *y* ≤6 arbitrary units. For specific details about the initial conditions, see Section S1.4.

### Source code

The source code for both the CPM and phase-field models are publicly available at https://github.com/DominicDevlin/Cell-differentiation-and-dewetting-drive-gut-villus-elongation.git

