## Supplementary Text and Figures for "Coupling cell differentiation to dewetting can explain villus elongation"

### Supplementary Information for *Coupling cell differentiation to dewetting can explain villus elongation*

### S1 Supplementary Text

#### S1.1 Supplementary Methods for the CPM

##### Tissue viscosity and surface tensions

In this section, we describe surface tensions in the CPM. Surface tensions are determined by the adhesion energies described in Appendix 4. Specifically, the surface tension between cell types (or medium) $\tau$ and $\tau'$, $\gamma_{\tau\tau'}$, is given by:

$$\gamma_{\tau\tau'} = J_{\tau\tau'} - \frac{J_{\tau\tau} + J_{\tau'\tau'}}{2}. \tag{S1}$$

For all simulations, cell-to-medium adhesion energies are equal, i.e., $J_{LM} = J_{SM} = J_M$ (which prevents the medium pushing on cells), and the medium self-adhesion, i.e., $J_{MM} = 0$. Thus,

$$\begin{aligned} \gamma_{LM} &= J_M - \frac{J_{LL}}{2} \\ \gamma_{SM} &= J_M - \frac{J_{SS}}{2} \\ \gamma_{SL} &= J_{SL} - \frac{J_{LL} + J_{SS}}{2}. \end{aligned} \tag{S2}$$

For simulations presented in the main text, Equation S2 yields $\gamma_{LM} = 2.33$, $\gamma_{SM} = 0.17$ and $\gamma_{LM} = 2.33$ (see Appendix 4).

Tissue viscosities in the CPM are determined by tension at cell boundaries. The only term in the Hamiltonian that affects tension at cell boundaries is adhesion energies. Since S-cells and L-cells form distinct clusters in our model, we can assume that self-state adhesion energies determine tissue viscosity, i.e., $J_{\tau\tau}$ (where $\tau \in \{\mathrm{L}, \mathrm{S}\}$). The chosen values of self-state adhesion energies in the main text, $J_{LL} = 0.66$ and $J_{SS} = 5.25$, correspond to fluid-like and solid-like behaviour, respectively.

In the subsequent section, we test the effect of varying surface tension on elongation length in the CPM. To vary surface tensions, we vary the adhesion energies $J_M$, $J_{SS}$ and $J_{SL}$ while keeping $J_{LL}$ constant. These adhesion energies are determined by the following equations derived from Eq. S2:

$$\begin{aligned} J_M &= \gamma_{LM} + J_{LL}/2 \\ J_{SS} &= 2\gamma_{LM} - 2\gamma_{SM} + J_{LL} \\ J_{SL} &= \gamma_{SL} + \gamma_{LM} - \gamma_{SM} + J_{LL}, \end{aligned}$$

These equations show that varying surface tension requires a change in either $J_{LL}$ or $J_{SS}$, thereby altering the L-tissue or S-tissue viscosity. We decided to keep the L-tissue viscosity constant because L-tissue viscosity plays a much larger role in elongation than S-tissue viscosity, as long as S-tissue is sufficiently solid.

**Introducing epithelial cells into the CPM** We adjusted our model to include an epithelial cell layer by replacing the passive medium with a population of hundreds of individual epithelial cells, hereafter referred to as E-cells (Fig. S7). We note that this model is more restrictive on mesenchymal cell motion than the epithelial layer in the real intestine, due to differences in the thickness of the epithelial cell layer. Specifically, the epithelial layer is only one or a few cell layers thick in the intestine, whereas it is at least ten layers thick in our model [9, 18]. All

E-cells are constrained to the same area as L-cells and S-cells, which is 80 pixels. Adhesion energies are determined as follows. We set $J_{LE} = J_{SE} = J_E$ and substitute into Equation S1 to get:

$$\begin{aligned} J_E &= \gamma_{LE} + J_{LL}/2 + J_{EE}/2 \\ J_{SS} &= 2\gamma_{LE} - 2\gamma_{SE} + J_{LL} \\ J_{SL} &= \gamma_{SL} + \gamma_{LE} - \gamma_{SE} + J_{LL}. \end{aligned}$$

Here, $J_{EE}$ is the intercellular adhesion energy between E-cells, $\gamma_{LE}$ denotes the surface tension at the interface between L-cells and E-cells, and $\gamma_{SE}$ denotes the surface tension at the interface between S-cells and E-cells. To change the viscosity of E-cells, we change $J_{EE}$, with increases in $J_{EE}$ making the epithelial tissue more viscous.

#### S1.2 Analysis of the dewet-differentiation mechanism in the CPM

To examine the robustness of the dewet-differentiation mechanism in the CPM, we asked how changes in surface tension affect the elongation length. To answer this, we conducted a parameter sweep of $\gamma_{SL}$, the surface tension between S-cells and L-cells, and $\gamma_{LM}$, the surface tension between L-cells and the medium (see Text S1 for surface tension derivations in the CPM). We set $\gamma_{SE} = 0.17$ for all simulations, which is small enough to prevent S-tissue from circularising. For each set of distinct parameters, we tracked (i) the elongation length achieved at the end of development, and (ii) the contact angle over development (Appendix 4.4), both averaged over 120 developmental replicates of the evolved morphology used in the main text. We ran simulations for 40,000 MCS, which we deemed sufficiently long for all L-cells to differentiate. We prematurely stop simulations if the interface size between L-cells and S-cells is zero (i.e., L-cells and S-cells separate) because the mesenchyme must remain intact to generate proper intestinal villi during real morphogenesis [18]. The results show that the elongation length strongly depends on both $\gamma_{SL}$ and $\gamma_{LM}$, which we arbitrarily separate into three outcomes. First, when $\gamma_{SL} > \gamma_{LM}$, elongation does not occur (Fig. S1A). Elongation does not occur because L-cells and S-cells separate (Fig. S1C), supported by a rapid rise in contact angles to $\pi$ radians at the start of development (Fig. S1B). Second, when $\gamma_{SL} \approx \gamma_{LM}$, extensive elongation occurs, resulting in the formation of finger-like villi (Fig. S1AD). The contact angles for villi that undergo extensive elongation remain high over development but do not reach $\pi$ radians, indicating that dewetting is occurring (Fig. S1B). Third, when $\gamma_{SL} < \gamma_{LM}$, elongation lengths decline, with the extent of elongation decreasing as the difference between these two surface tensions increases (Fig. S1AIJ). Similarly, contact angles decline with the extent of elongation (Fig. S1B), indicating that dewetting is essential for the formation of finger-like villi. These results show that elongation via the dewet-differentiation mechanism is robust to changes in surface tensions so long as there is a balance between $\gamma_{SL}$ and $\gamma_{LM}$, in agreement with the results from our continuum model.

#### S1.3 Erroneous dewetting dynamics at high surface tensions in the CPM

A discrepancy between the continuum model results and CPM results is lower elongation lengths at high surface tensions in the CPM (top-right corner of Fig. S1A). Specifically, when $\gamma_{LM}$ and $\gamma_{SL}$ are high, the force driving dewetting is higher, which we expected would promote elongation. We hypothesised that this reduction may be due to dewetting behaviour in the CPM. To test this hypothesis, we recorded contact angles across the simulations in the parameter sweep of surface tensions shown in Fig. S1A (since contact angles measure wetting/dewetting). We plotted the average contact angle between 8,000 to 10,000 MCS against the final elongation for each simulation. We found that contact angles paradoxically decreased when surface tensions are

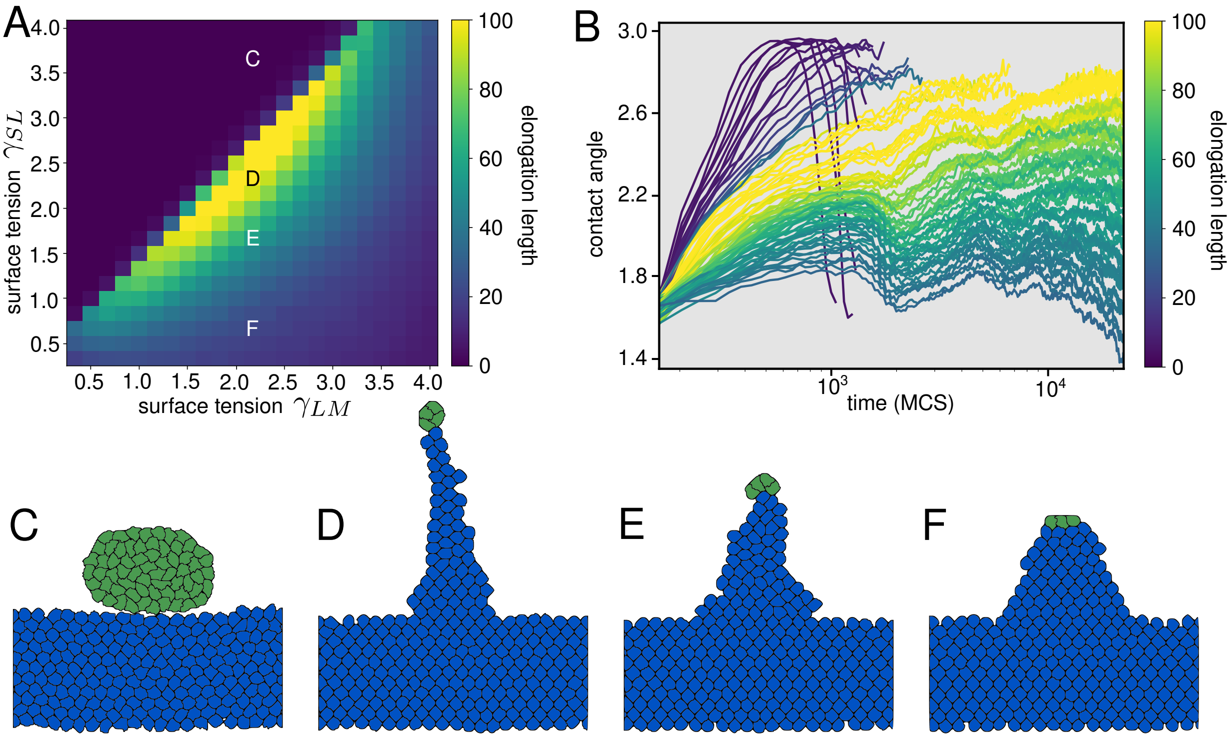


Figure S1: **The effect of surface tension on villus elongation. A** Phase diagram of the elongation length as a function of surface tension at the interface between L-cells and the medium ($\gamma_{LM}$), and surface tension at the interface between S-cells and L-cells ($\gamma_{SL}$). Each data point is the average elongation length over 120 developmental replicates. **B** Contact angles over logarithmic time for a sample of the data points in (A). Each line is colour-coded by elongation length. Each line averages over the 120 developmental replicates, using the same surface tension simulations as in (A). The rapidly increasing and decreasing contact angles correspond to morphologies where the L-cells and S-cells separate. **C,D,E,F** Snapshots of endpoint morphologies using $\gamma_{LM} = 2.2$ and four different values of $\gamma_{SL}$: (C) 3.7, (D) 2.3, (E) 1.7 and (F) 0.7.

too high (Fig. S2A), contrary to predictions based on classical wetting theory [16]. Specifically, the results show a non-linear relationship between the contact angle and surface tensions: the contact angles increase when $\gamma_{LM}$ and $\gamma_{SL}$ are less than $\sim 2.5$, but then decrease as $\gamma_{LM}$ and $\gamma_{SL}$ increase above $\sim 2.5$ (Fig. S2A). This result suggests that dewetting of L-cells from S-cells is impaired when surface tensions get too high in the CPM.

To test why dewetting is impaired at high surface tensions, we simplified our model to simulations of L-cell clusters surrounded by the medium. There are no S-cells. We simulated the circularisation of these L-cell clusters starting from a semicircular shape. Physically, the time required to circularise should decrease as the energy arising from surface tension increases. We simulated the development of these clusters for 15,000 MCS. Fig. S2C-E shows snapshots of L-cell clusters after 15,000 MCS with low $\gamma_{LM}$ (C), intermediate $\gamma_{LM}$ (D) and high $\gamma_{LM}$ (E). The snapshots show that the cluster with intermediate $\gamma_{LM}$ exhibits the most circularity, whereas the cluster with high $\gamma_{LM}$ exhibits a polygonal shape with flat instead of curved surfaces at the interface with the medium. To confirm this observation, we measured the circularity of the L-cell cluster 15,000 MCS. The circularity, denoted by $z$, is defined by the following equation:

$$\langle |r_c - r(\theta)| \rangle_\theta, \tag{S3}$$

where $r_c$ is the hypothetical radius of the morphology if it were perfectly circular, and $r(\theta)$ is the maximum distance from the centre of mass of the morphology to any pixel in the direction specified by angle $\theta$ (one pixel corresponds to one unit mass). The notation $\langle ... \rangle_\theta$ indicates an average over angles $\theta$ (we divide $\theta$ into 360 elements for computation). The results show that $z$ decreases with $\gamma_{LM}$ until a minimum is reached at $\gamma_{LM} \approx 2.3$ (Fig. S2B). When $\gamma_{LM}$ increases above 2.3, $z$ also increases until reaching a plateau. This value of 2.3 is in agreement with the value of 2.5 deduced from Fig. S2A. From these results, we deduce that at high surface

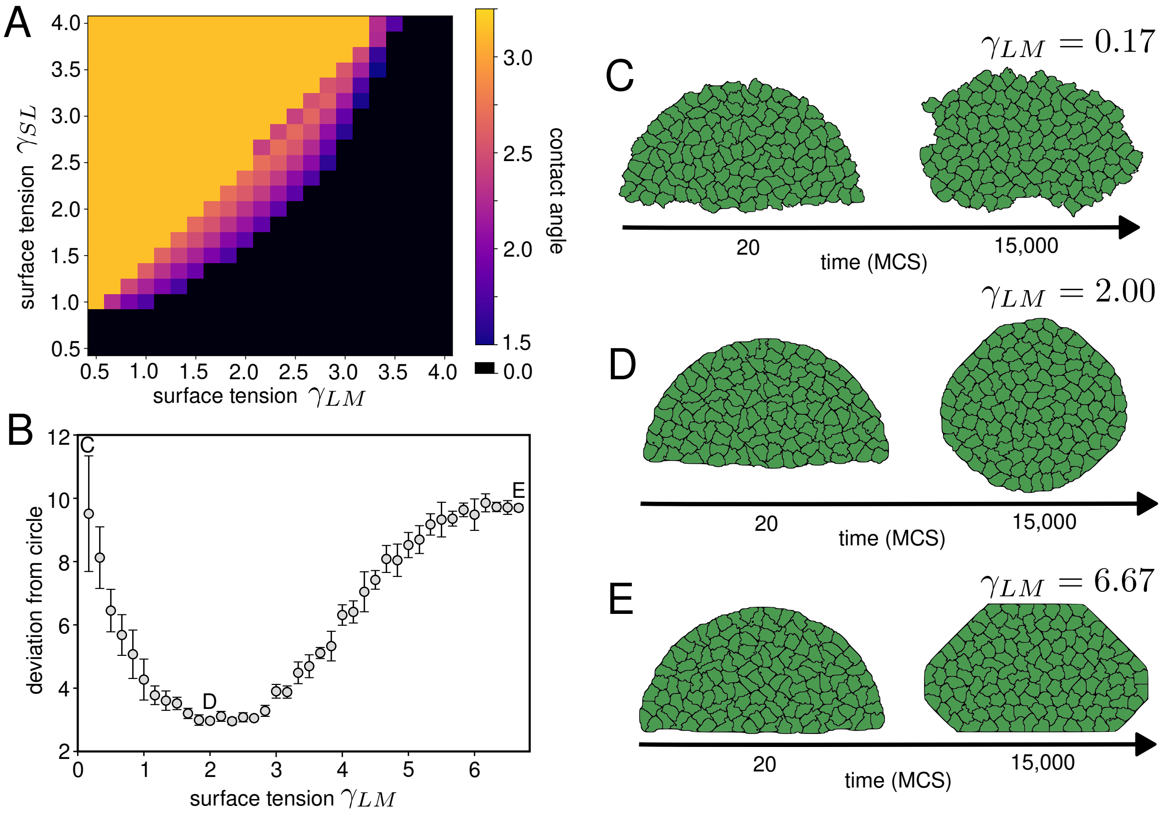


Figure S2: **CPM dynamics at high surface tensions.** **A** Phase diagram of contact angles after 10,000 MCS as a function of $\gamma_{LM}$ and $\gamma_{SL}$ for the same set of simulations shown in Fig. S1A. We prescribe a contact angle of zero if all L-cells have differentiated, and a contact angle of $\pi$ if none have differentiated. This phase diagram indicates non-physical behaviour because contact angles decrease as surface tension increases along the diagonal. The colour indicates the contact angle as shown by the heatmap. The colours between 0.0 and 1.5 have been removed from the heatmap fro visibility. **B** The deviation of a morphology from a circle as a function of the surface tension at the LM-interface. Each data point is the average deviation of a circle between 8,000 and 10,000 MCS and averaged over 120 developmental replicates ($\pm$ standard deviation). Each replicate starts as a semicircle of cells surrounded by the medium. **C,D,E** Three example morphologies starting from the semicircular arrangement with (C) $\gamma_{LM} = 0.17$, (D) $\gamma_{LM} = 2.00$, and (E) $\gamma_{LM} = 6.67$. Each morphology is shown after 20 MCS and after 15,000 MCS.

tensions, the increased sensitivity to energy at cell boundaries causes the multicellular cluster to adopt a polygonal configuration that minimises local interface length. Polygonal configuration minimises local interface length because the grid comprises discrete pixels, i.e., it is not continuous. Thus, the non-linear relationship between contact angle and surface tension is due to the discretisation of the CPM grid into pixels, rather than a physical phenomenon. Thus, the CPM is inappropriate for examining the dewet-differentiation mechanism at high surface tensions without modifications to minimise the effects of grid discretisation.

#### S1.4 Supplementary Methods for the Continuum model

**Surface tension derivation.** The surface tension in a two-dimensional system represents the energetic cost of maintaining the interface per unit length [45]. Without external forces, tissues in the phase-field model minimise surface tension by adopting circular morphologies, consistent with the behaviour of real biological tissues [46]. We derive the surface tension for a binary interface between phases $i$ and $j$ through contributions from the gradient energy density and bulk potential in Equation 5 (Appendix 5). That is, we derive the surface tension by assuming that phases shift strictly along the boundary of the ternary plot shown in Fig. S3. This assumption holds in our continuum model everywhere except triple points, where the three phases meet. To derive the energetic cost per unit length, we consider an equilibrium profile $c(x)$ across a flat interface between phases $i$ and $j$ normal to the $x$-axis, with $c_i = c$ and

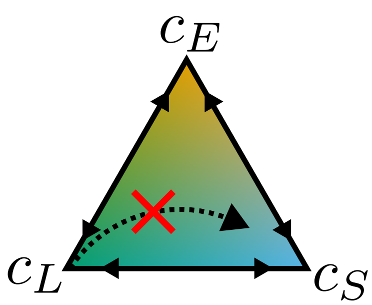


Figure S3: **Assumption for surface tension derivation illustrated on a ternary plot.** The path along the two-sided arrows indicates binary interfaces from which we derive surface tension. The dashed arrow indicates a path not accounted for in the surface tension derivation (although these paths are only found at triple points).

$c_j = 1 - c$. The gradient term in the functional (Eq. 7) has the form

$$\epsilon^2 \kappa_{ij} \left| c_i \nabla c_j - c_j \nabla c_i \right|^2.$$

Using $c_j = 1 - c_i$ and $\nabla c_j = -\nabla c_i$, we obtain

$$c_i \nabla c_j - c_j \nabla c_i \;=\; c(-\nabla c) - (1-c)\nabla c \;=\; -\nabla c,$$

so that, along the $x$-axis the energy is:

$$\epsilon^2 \kappa_{ij} \left| c_i \nabla c_j - c_j \nabla c_i \right|^2 = \epsilon^2 \kappa_{ij} \, (\partial_x c)^2,$$

where $\partial_x c$ is the spatial derivative of $c$ with respect to $x$. In our model, the local free-energy density arising from the bulk potential between two phases is (Eq. 8):

$$\kappa_{ij} c_i^2 c_j^2 = \kappa_{ij} \, c^2 (1-c)^2.$$

Thus, the total energy along the one-dimensional profile (i.e., the surface tension) is:

$$\gamma_{ij} \;=\; \int_{-\infty}^{\infty} \left[ \epsilon^2 \kappa_{ij} \, (\partial_x c)^2 \;+\; \kappa_{ij} \, c^2 (1-c)^2 \right] \mathrm{d}x. \tag{S4}$$

Since the profile is at equilibrium, the gradient energy density and the bulk potential balance, giving:

$$\epsilon^2 \kappa_{ij} \, (\partial_x c)^2 \;=\; \kappa_{ij} \, c^2 (1-c)^2 \;\Rightarrow\; \partial_x c \;=\; \frac{c(1-c)}{\epsilon}. \tag{S5}$$

We then change the variables in Eq. S4 from $dx$ to $dc$ by substituting for $dx$ via Eq. S5 and using the boundary conditions $c(-\infty) = 0$, $c(+\infty) = 1$, giving:

$$\gamma_{ij} = 2 \int_{-\infty}^{\infty} \epsilon^2 \kappa_{ij} \, (\partial_x c)^2 \, \mathrm{d}x = 2\epsilon\kappa_{ij} \int_0^1 c(1-c) \, \mathrm{d}c = \frac{\epsilon \, \kappa_{ij}}{3}.$$

Hence,

$$\boxed{\gamma_{ij} = \frac{\epsilon \, \kappa_{ij}}{3}} \tag{S6}$$

for each pair $i, j \in \{S, L, E\}$. Since we used $\epsilon = 0.02$ for all simulations, $\gamma_{ij} = 0.02\kappa_{ij}/3$. The corresponding equilibrium concentration profile along the $x$-axis follows from an integration of Eq. S5:

$$\partial_x c = \frac{c(1-c)}{\epsilon} \quad \Rightarrow \quad c(x) = \frac{1}{1 + \exp(-x/\epsilon)}. \tag{S7}$$

Equation S7 shows that, at equilibrium, the concentration profile is independent of $\kappa_{ij}$, which means different kinds of interfaces have the same width independent of the surface tension at each interface. From Equation S7, we can determine the width of the interface at equilibrium. For instance, between $c = 0.05$ and $c = 0.95$ with $\epsilon = 0.02$ the thickness is $\approx 0.12$.

**Initial Conditions.** For simulations presented in the main text, we formulate an initial condition of S-tissue and L-tissue that resembles an intestinal villus after the initial clustering of L-tissue [13]. For the S-tissue, $c_S$, the initial condition is a flat sheet that wraps around the periodic boundaries horizontally, as follows:

$$c_S = \frac{1}{2}\left[1 - \tanh\left(\frac{|\,y - y_c\,| - H}{\delta_b}\right)\right],$$

where $H = 0.7$ is half the length of the tissue along the $y$-axis, $y_c = 1.0$ is the centre of the tissue along the $y$-axis and $\delta_b = 0.03$ determines the initial tissue interface thickness. For the L-tissue, $c_L$, the initial condition is a semicircle that rests on top of the S-tissue with radius $R = 0.8$. To create the semicircle, we first define a smooth disk function $U(x, y)$, where $c_L \to 1$ inside the disk and $c_L \to 0$ outside. This is achieved using the hyperbolic tangent function as follows:

$$U(x, y) = \frac{1}{2}\left[1 - \tanh\left(\frac{r(x, y) - R}{\delta_b}\right)\right],$$

where $r(x, y) = \sqrt{(x - x_0)^2 + (y - y_0)^2}$ is the distance from the centre. We set $x_0 = 2$, which is the horizontal centre of the domain, and $y_0 = y_c + H$, so that the centre is in the same location vertically as the S-tissue interface. Next, we define a mask $M(y)$ to smoothly separate the upper and lower halves of the circle, defined as:

$$M(y) = \frac{1}{2}\left[1 + \tanh\left(\frac{y - y_0}{\delta_b}\right)\right],$$

The semicircular initial condition is obtained by multiplying the smooth disk function with the upper-half mask, i.e., $c_L = U(x, y)M(y)$. Until $t = 0.2$, we allow the system to relax from this initial condition by setting $V_S(t) = v_S(t)$ and $V_L(t) = v_L(t)$ for all $t$.

For the results shown in Fig. S9, we simulate the entire process of intestinal villus morphogenesis by initialising L-tissue as a flat sheet instead of a semicircle, which then forms clusters and develops into elongated villi. For this initial condition, we define a smooth horizontal band function $X(x)$ and a smooth vertical band function $Y(y)$, where $c_L \to 1$ inside the bands and $c_L \to 0$ outside. This is achieved using hyperbolic tangent functions as follows:

$$X(x) = \tfrac{1}{2}\big(1 + \tanh([x - x_{\text{start}}]/\delta_b)\big) - \tfrac{1}{2}\big(1 + \tanh([x - x_{\text{end}}]/\delta_b)\big)$$
$$Y(y) = \tfrac{1}{2}\big(1 + \tanh([y - y_{\text{start}}]/\delta_b)\big) - \tfrac{1}{2}\big(1 + \tanh([y - y_{\text{end}}]/\delta_b)\big),$$

where $X(x) > 0.5$ between $x = x_{\text{start}}$ and $x = x_{\text{end}}$, and $Y(y) > 0.5$ between $y = y_{\text{start}}$ and $y = y_{\text{end}}$. We set:

$$y_{\text{start}} = y_0, \qquad y_{\text{end}} = y_{\text{start}} + \mathrm{H}, \qquad x_{\text{start}} = x_0 - \frac{\mathrm{W}}{2}, \qquad x_{\text{end}} = x_0 + \frac{\mathrm{W}}{2},$$

where $x_0$ and $y_0$ are the same as the previous initial condition, W is the width of the L-tissue sheet and H is the height of the L-tissue sheet. We set $\mathrm{W} = 3$ and $\mathrm{H} = 0.36$, so that the L-tissue sheet is much wider than it is tall. The initial condition is obtained by multiplying the smooth band functions together, i.e., $c_L = X(x)Y(y)$. For this initial condition, we extended the time at which $D = 0$ to $t = 100$ (it was until $t = 10$ for simulations in the main text), so that the L-tissue has sufficient time to form clusters before differentiating.

#### S2 Supplementary Figures

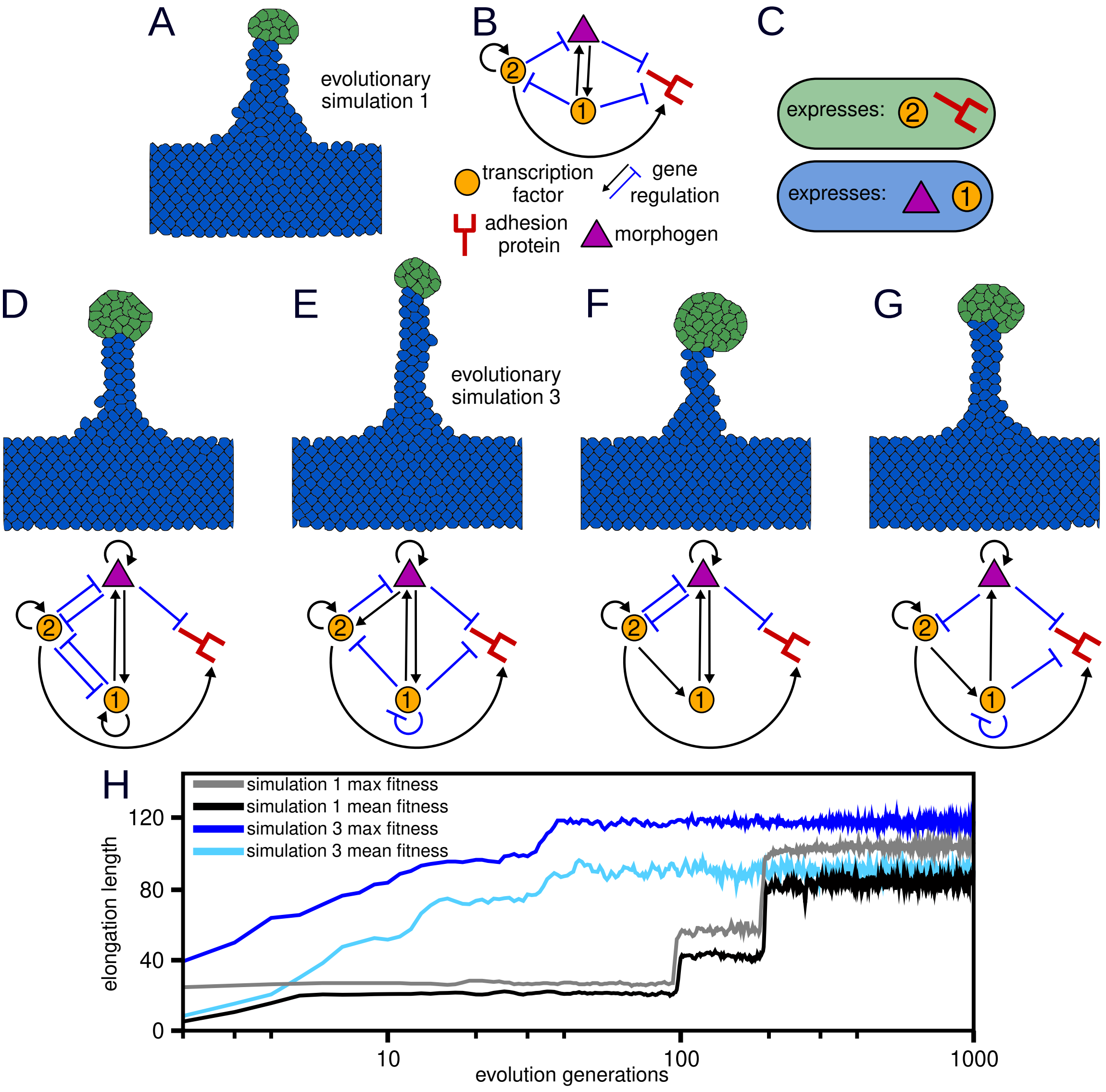


Figure S4: **Evolution of elongation in the CPM. A** Evolved morphology developed for 20,000 MCS (half the total simulation time). This evolved morphology is the one shown in the main text from evolutionary simulation 1. **B** Evolved GRN used to simulate the morphology shown in (A). The schematic shows three transcription factors (TFs; circles and triangle) and one protein (red stick) that determines the cell type: L-cell or S-cell. The triangle represents a membrane-permeable TF (the morphogen) that diffuses between cells. Arrows indicate the regulation of gene expression by TFs (black for activation, light blue for inhibition). **C** Diagram showing the proteins expressed by L-cells (green box) and S-cells (blue box) for the evolved morphology shown in (A). **D,E,F,G** Evolved morphologies developed for 20,000 MCS (half the total simulation time) using the most common GRN observed in the 1,000$^{th}$ generation from simulations 2-5. Below each morphology is the GRN logic for that morphology. Common patterns in GRNs include the self-activation of the morphogen, self-activation of TF-2, and the activation of the adhesion protein by TF-2. **H** Elongation length as a function of evolution generations for two separate simulations (generations are in log scale). The grey (simulation 1) and dark blue (simulation 3) lines plot the elongation length of the morphology with the highest elongation length in each generation from the two simulations. The black (simulation 1) and light blue (simulation 3) lines correspond to the average elongation length in each generation from the two simulations.

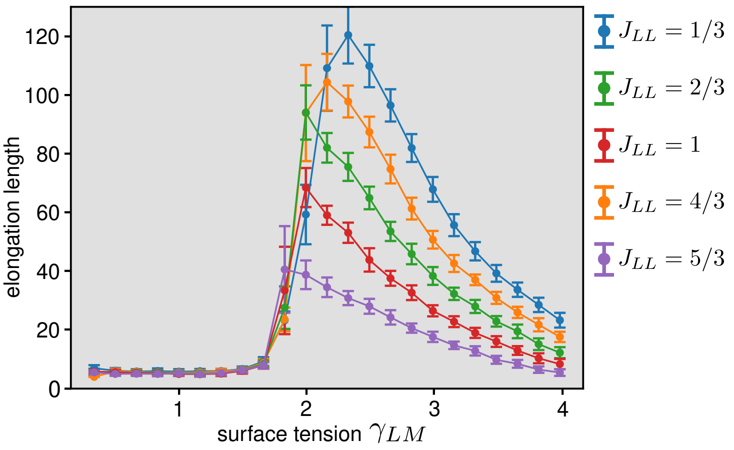


Figure S5: **The effect of L-tissue viscosity on elongation length in the CPM.** The plot shows the elongation length as a function of $\gamma_{LM}$ with $\gamma_{SL} = 2.33$ for the evolved morphology shown in the main text. Each line plots a different value of $J_{LL}$, the adhesion energy between L-cells. Higher $J_{LL}$ corresponds to a more viscous L-tissue. Each data point is the average elongation length ($\pm$ standard deviation), averaged over 120 developmental replicates. As viscosity increases, there is a decline in elongation length at each $\gamma_{LM}$.

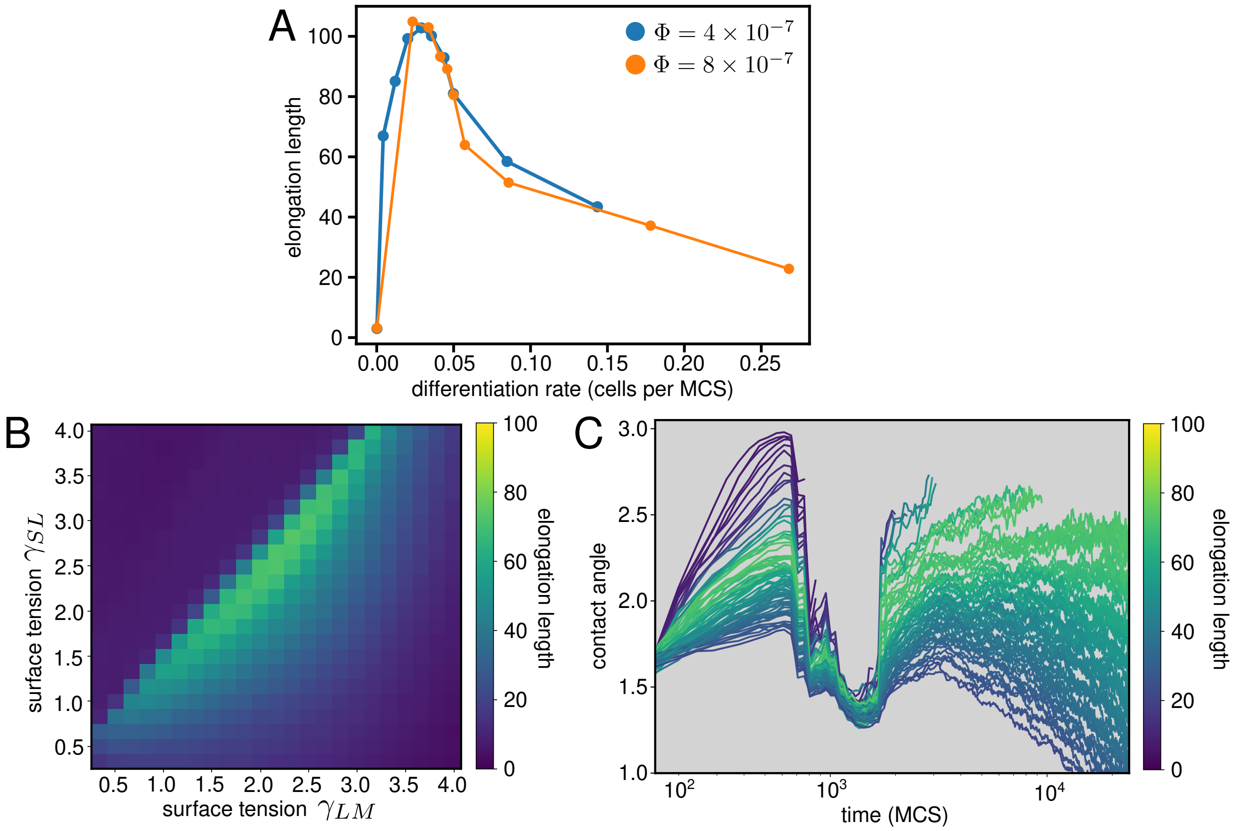


Figure S6: **The effect of differentiation rate on elongation length in the CPM. A** Elongation length as a function of the differentiation rate. The differentiation rate is the number of L-cells that differentiate per MCS, measured until half the total simulated time (simulations stop when all L-cells differentiate). The differentiation rate and elongation lengths are averaged over 120 developmental replicates. We tune the differentiation rate via the morphogen production rate, $\omega$ and the diffusion rate, $\Phi$ (see Eq. 4). The blue line shows the elongation lengths with $\Phi = 4 \times 10^{-7}$ for $\omega \in \{0, 0.002, 0.00225, 0.0025, 0.00275, 0.003, 0.0035, 0.004, 0.005, 0.006\}$. The orange line shows the elongation lengths with $\Phi = 8 \times 10^{-}7$ for the same ten values of $\omega$. **B** Elongation length as a function of $\gamma_{LM}$ and $\gamma_{SL}$. Each data point is the average elongation length across 120 developmental replicates, with each replicate lasting up to 40,000 MCS. For all simulations, $\omega = 0.006$ and $\Phi = 4 \times 10^{-7}$, with these parameters corresponding to a higher differentiation rate than all other simulations (see Appendix 4). This high differentiation rate causes a global reduction in elongation length compared to Fig. S1A. **C** Contact angles over logarithmic time for a sample of the data points in (B). Each line is colour-coded by elongation length. For each line, each time point averages over 120 developmental replicates. The significant decrease in contact angles at $\sim 10^3$ MCS is caused by the first layer of L-cells in the semicircular cluster differentiating all at once because of the high differentiation rate.

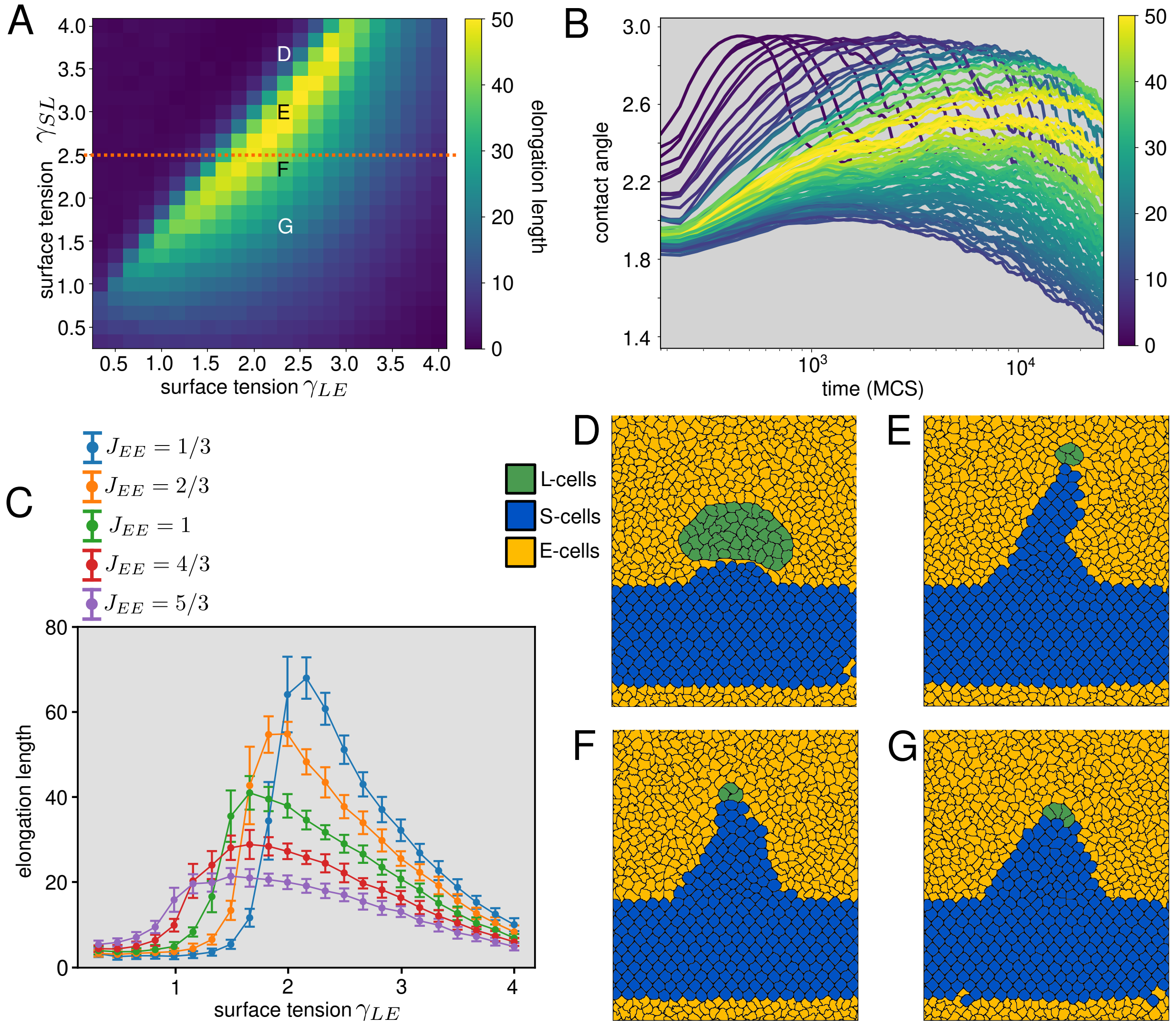


Figure S7: **Villus elongation in the CPM with an epithelial cell cage**. To assess the impact of the epithelial cage on villus elongation, we replaced the medium with a layer of epithelial cells (Text S1.1). We explored how variations in surface tensions influence elongation relative to our previous simulations using the medium. We performed a parameter sweep of $\gamma_{LE}$ and $\gamma_{SL}$, tracking the elongation length achieved, averaging over 120 developmental replicates. We used the same evolved morphology as used in the main text. We ran simulations until 40,000 MCS or until the L-cells and S-cells separated. **A** Phase diagram of the elongation length as a function of $\gamma_{LE}$ and $\gamma_{SL}$. Each data point is the elongation length averaged across 120 developmental replicates, with each replicate lasting up to 40,000 MCS. $\gamma_{SE} = 1/12$ for all simulations. **B** Contact angles over time for the same data points shown in (A), colour-coded by elongation length, with a logarithmic time scale. Each line is averaged over 120 developmental replicates. **C** Elongation length in the presence of the epithelial cage as a function of $\gamma_{LE}$ for $\gamma_{SL} = 2.5$. The orange, green, purple and red lines show results for different values of $J_{EE}$, with higher $J_{EE}$ corresponding to higher viscosity of the epithelial cage. The blue line shows elongation with the medium instead of epithelial cells as a function of $\gamma_{LM}$. Error bars are standard deviations on the length elongated across 120 developmental replicates. **D,E,F,G** Snapshots of endpoint morphologies in the epithelial layer model with $\gamma_{LE} = 2.33$ and $\gamma_{SE} = 1/12$ for four different values of $\gamma_{SL}$: (D) 3.67, (E) 3.0, (F) 2.33 and (G) 1.67.

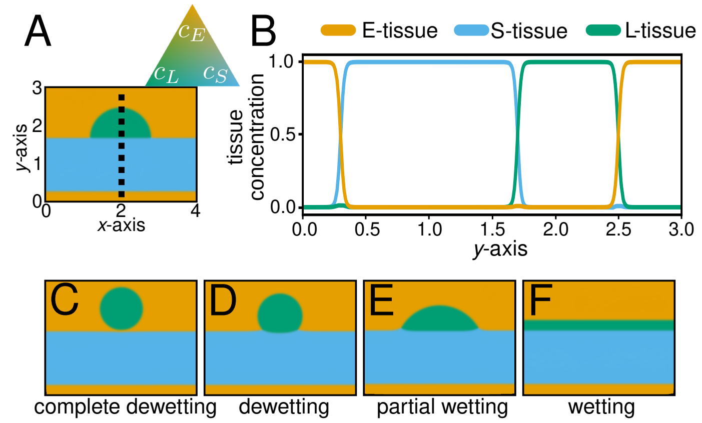


Figure S8: **Continuum model of villus elongation.** **A** The initial tissue configuration in the continuum modem for simulations in the main text. The colour at each point in the snapshot represents the local concentrations of S-tissue ($c_S$), L-tissue ($c_L$), and epithelial tissue ($c_E$). Snapshots are coloured by the concentration of $c_S$, $c_L$, and $c_E$ as indicated by the ternary plot with vertices of $c_S = c_L = c_E = 1$. We show a subsection of the domain ($0 \leq y \leq 3$ instead of $0 \leq y \leq 6$) for improved visibility. **B** Concentrations of $c_S$, $c_L$ and $c_E$ along a cross-section of the snapshot in (A) depicted by the dashed black line. **C-F** Four snapshots of tissue configurations after simulating our model until $t = 500$ when there is no differentiation, with each snapshot showing a simulation with different surface tensions: (C) complete dewetting ($\gamma_{SL} = 0.14, \gamma_{LE} = 0.06$), (D) dewetting ($\gamma_{SL} = 0.14, \gamma_{LE} = 0.10$), (E) partial wetting ($\gamma_{SL} = 0.02, \gamma_{LE} = 0.06$) and (F) wetting ($\gamma_{SL} = 0.02, \gamma_{LE} = 0.02$); $\gamma_{SE} = 0.06$ for all simulations. The snapshot in (F) was taken at $t = 3000$, as it required longer to equilibrate.

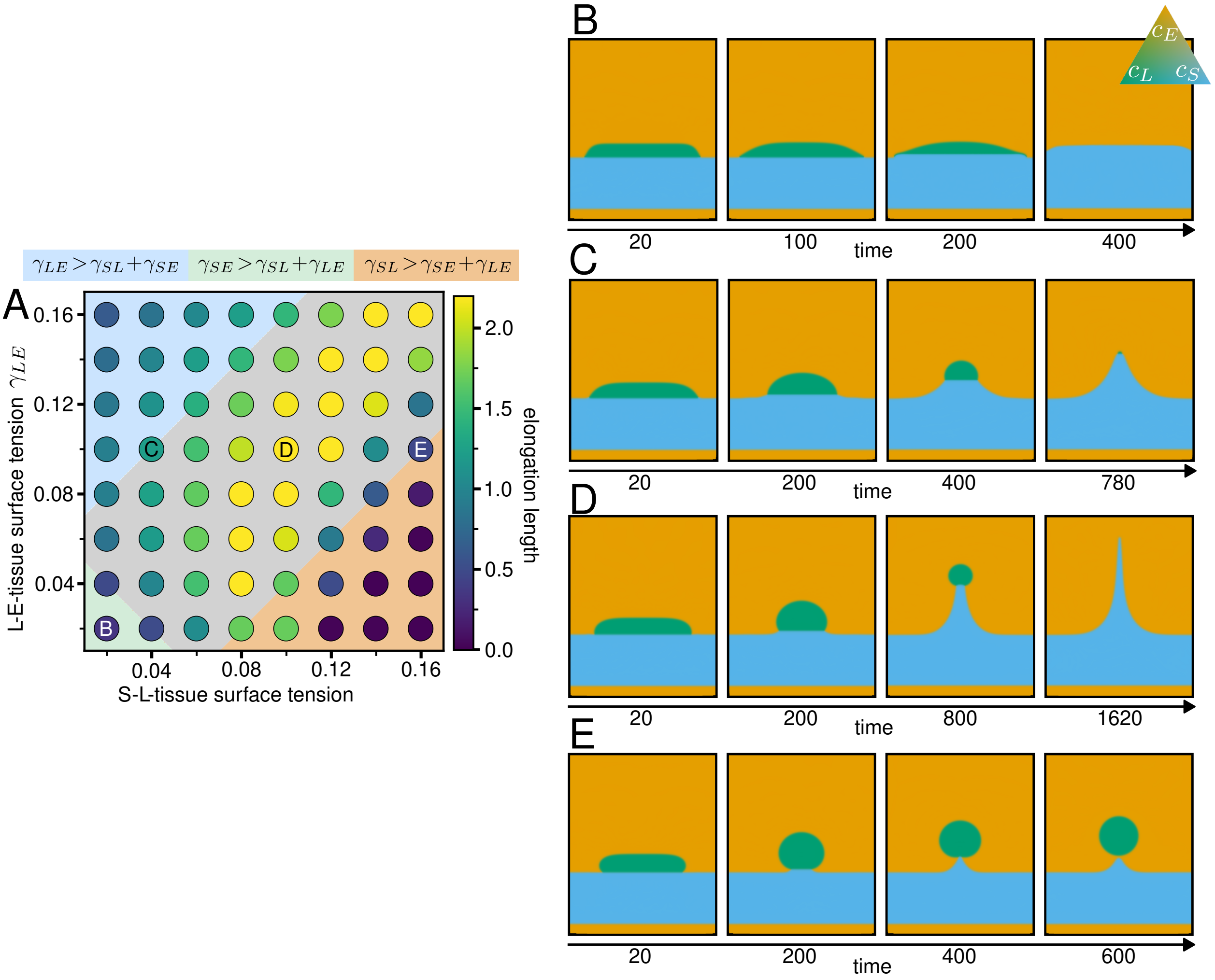


Figure S9: **Villus elongation starting from a flat L-tissue sheet.** For simulations shown in this figure, we use the second initial condition outlined in Text S1.4. **A** Phase diagram showing simulation-end elongation length as a function of $\gamma_{SL}$ and $\gamma_{LE}$. The background colour of the phase diagram indicates the spreading regime, as indicated on top (blue is where S-tissue spreads ($\mathbf{S}_S > 0$), green is where L-tissue spreads ($\mathbf{S}_L > 0$), and orange is where E-tissue spreads ($\mathbf{S}_E > 0$)). Grey is the non-spreading regime ($\mathbf{S}_i < 0$ for $i = S, L, E$). **B,C,D,E** Snapshots of elongating villi at consecutive time-points in their respective simulations, with the time-points of each shown below the snapshot. Snapshots are coloured by the concentration of $c_S$, $c_L$, and $c_E$ as indicated by the ternary plot with vertices of $c_S = c_L = c_E = 1$. The values of $\gamma_{SL}$ and $\gamma_{LE}$ for each simulation are shown in (A). For (B), (C), and (D), the final snapshot is taken approximately when the L-tissue volume falls below 0.001. For (E), the final snapshot is taken approximately when L-tissue and S-tissue separate. $\gamma_{SE} = 0.06$, $D = 1$, $\eta_L = 1$ and $\eta = 100$ for all simulations.

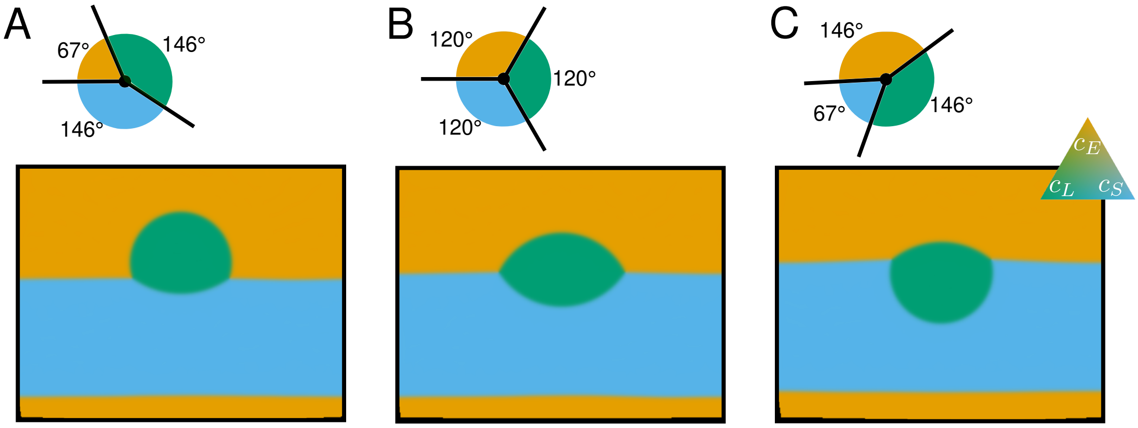


Figure S10: **Equilibrium contact angles in the continuum model coincide with predicted equilibrium contact angles.** To assess whether contact angles in our model recover those predicted from classical wetting theory, we simulated our model with $D = 0$, $\eta_S = 1$, and $\eta_L = 1$ until $t = 500$ (a time sufficient for the configurations to reach equilibrium). The three snapshots in this figure show simulations at time $t = 500$. The surface tensions used are: **A** $\gamma_{SE} = 0.06$, $\gamma_{SL} = 0.10$ and $\gamma_{LE} = 0.06$ **B** $\gamma_{SE} = 0.06$, $\gamma_{SL} = 0.06$ and $\gamma_{LE} = 0.06$ **C** $\gamma_{SE} = 0.06$, $\gamma_{SL} = 0.06$ and $\gamma_{LE} = 0.10$. Snapshots are coloured by the concentration of $c_S$, $c_L$, and $c_E$ as indicated by the ternary plot with vertices of $c_S = c_L = c_E = 1$. Above each snapshot, a schematic shows the predicted equilibrium contact angles at the triple point, calculated from surface tension values using the Neumann triangle relation. For the angle measured through phase $i$, i.e., between interfaces $i{-}j$ and $i{-}k$ with $\{i, j, k\} = \{L, S, E\}$, the formula is $\cos\theta_i = \frac{\gamma_{jk}^2 - \gamma_{ik}^2 - \gamma_{ij}^2}{2\,\gamma_{ij}\,\gamma_{ik}}$. Arc colours indicate the phase through which the angle is taken: green = L-tissue, blue = S-tissue, orange = E-tissue.

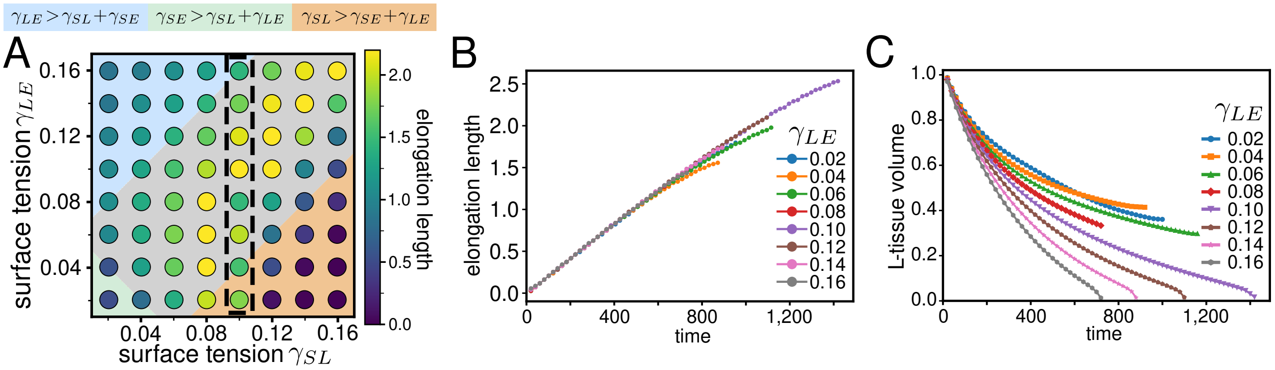


Figure S11: **Supplementary analysis of villus elongation in the continuum model.** **A** Phase diagram showing simulation-end elongation length (circle colour, see heatmap) as a function of $\gamma_{SL}$ and $\gamma_{LE}$ (replica of Fig. 3H). The background colour of the phase diagram indicates the spreading regime, as indicated on top (blue is where S-tissue spreads ($\mathbf{S}_S > 0$), green is where L-tissue spreads ($\mathbf{S}_L > 0$), and orange is where E-tissue spreads ($\mathbf{S}_E > 0$)). Grey is the non-spreading regime ($\mathbf{S}_i < 0$ for $i = S, L, E$). **B** Elongation length and **C** L-tissue volume as a function of time for different values of $\gamma_{LE}$. Each line corresponds to a simulation inside the dashed black box in (A). $\gamma_{SL} = 0.1$, $\gamma_{SE} = 0.06$, $\eta = 100$ and $D = 1$ for all simulations.

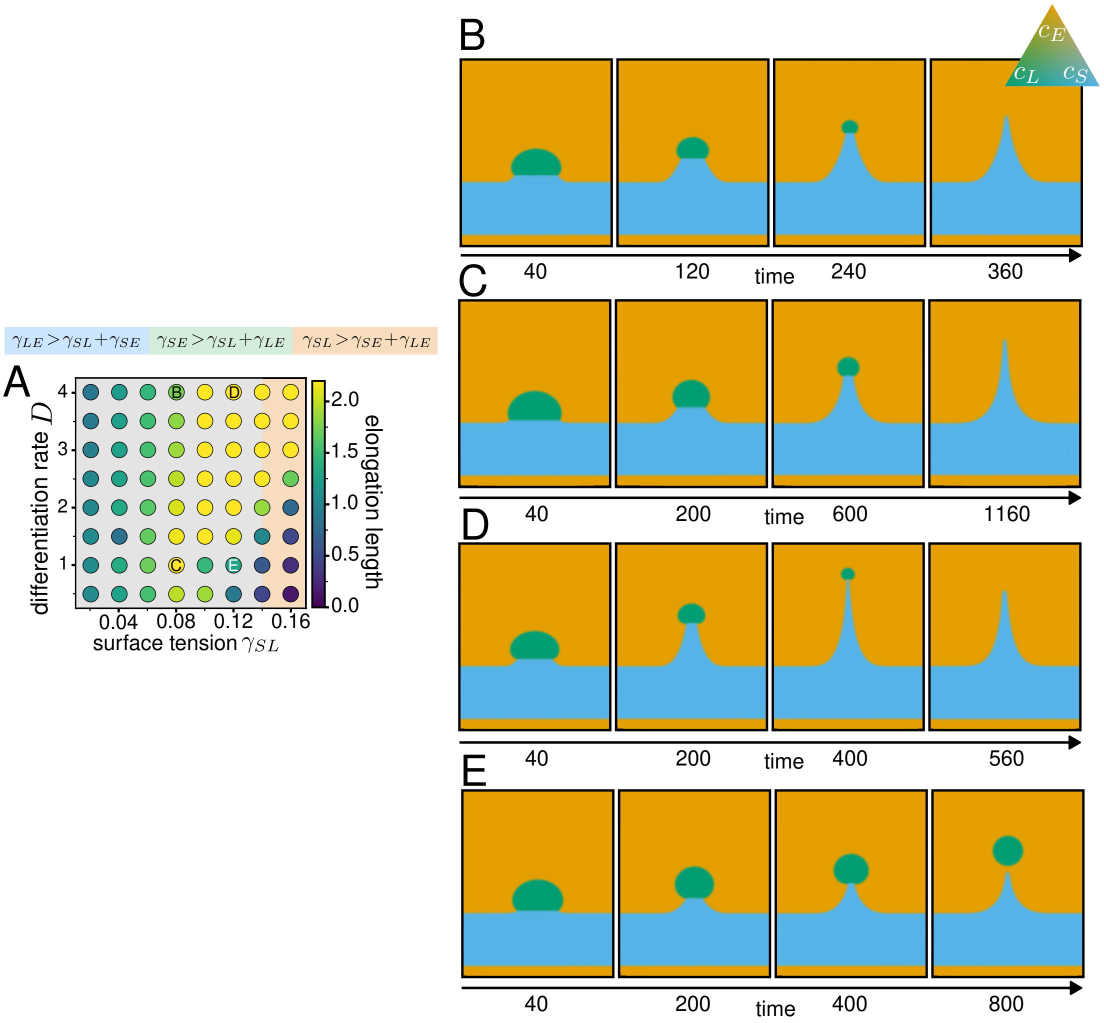


Figure S12: **The effect of the differentiation parameter $D$ on villus morphologies. A** Phase diagram showing simulation-end elongation length (circle colour, see heatmap) as a function of $\gamma_{SL}$ and $D$ (replica of Fig. 3K). The background colour of the phase diagram indicates the spreading regime, as indicated on top (blue is where S-tissue spreads ($\mathbf{S}_S > 0$), green is where L-tissue spreads ($\mathbf{S}_L > 0$), and orange is where E-tissue spreads ($\mathbf{S}_E > 0$)). Grey is the non-spreading regime ($\mathbf{S}_i < 0$ for $i = S, L, E$). **B,C,D,E** Snapshots of four elongating villi at consecutive time-points in a simulation, with each time-point shown below the snapshot. Snapshots are coloured by the concentration of $c_S$, $c_L$, and $c_E$ as indicated by the ternary plot with vertices of $c_S = c_L = c_E = 1$. $\gamma_{SE} = 0.06$, $\gamma_{LE} = 0.08$ and $\eta = 100$ for all simulations. See (A) for values of $D$ and $\gamma_{SL}$. The snapshots show that increasing $D$ can either decrease elongation length (B vs C) or increase elongation length (D vs E) depending on $\gamma_{SL}$.

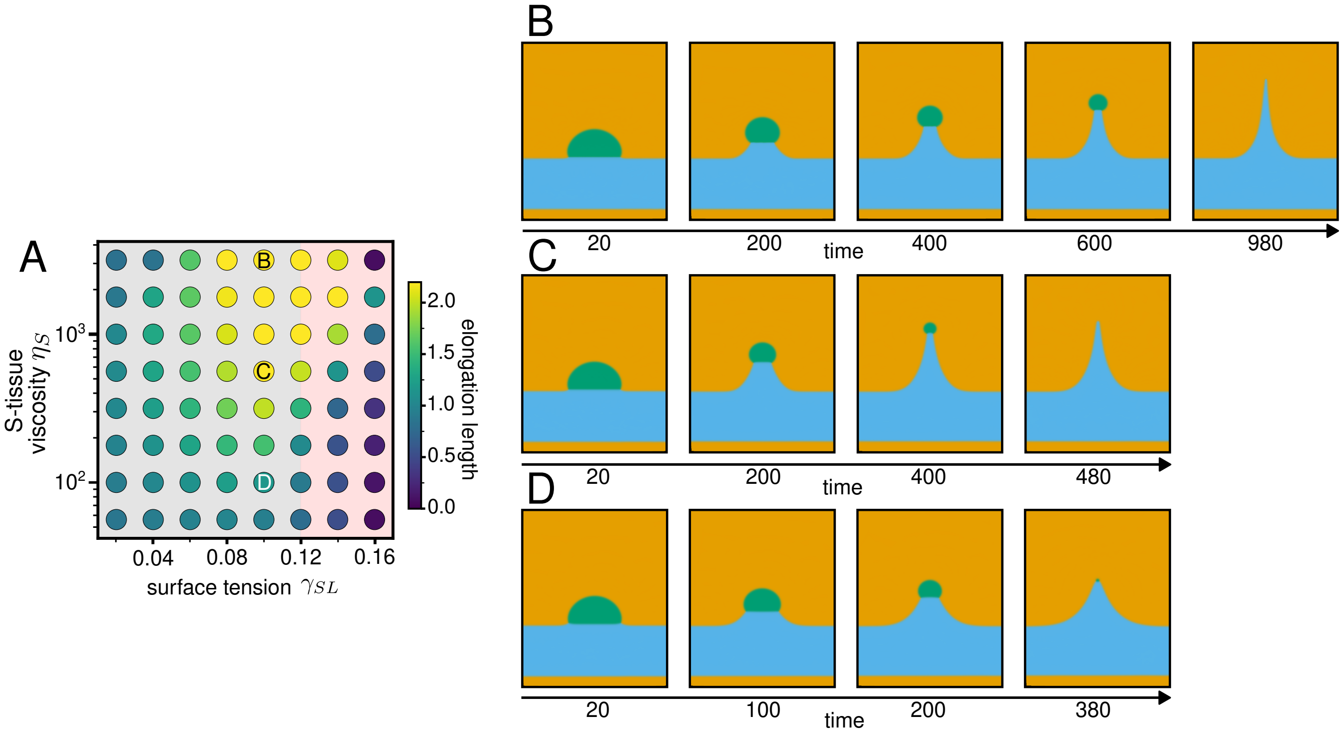


Figure S13: **The effect of relative viscosity $\eta$ on villus morphologies. A** Phase diagram showing simulation-end elongation length (circle colour, see heatmap) as a function of $\gamma_{SL}$ and $\eta$ (replica of Fig. 3N). The background colour of the phase diagram indicates the spreading regime, as indicated on top (blue is where S-tissue spreads ($\mathbf{S}_S > 0$), green is where L-tissue spreads ($\mathbf{S}_L > 0$), and orange is where E-tissue spreads ($\mathbf{S}_E > 0$)). Grey is the non-spreading regime ($\mathbf{S}_i < 0$ for $i = S, L, E$). **B,C,D** Snapshots of three elongating villi at consecutive time-points in a simulation, with each time-point shown below the snapshot. Snapshots are coloured by the concentration of $c_S$, $c_L$, and $c_E$ as indicated by the ternary plot with vertices of $c_S = c_L = c_E = 1$. $\gamma_{SE} = 0.06$, $\gamma_{LE} = 0.08$ and $D = 2$ for all simulations. See (A) for values of $\eta$ and $\gamma_{SL}$. The snapshots show that S-tissue resists being stretched into a finger-like morphology when $\eta$ is small.

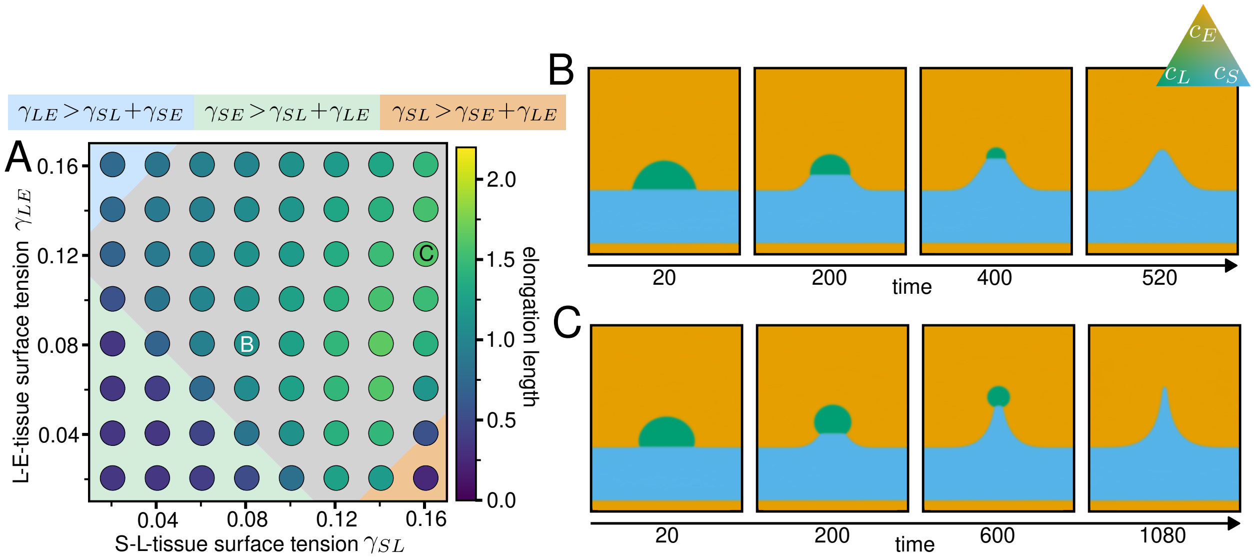


Figure S14: **The effect of higher SE-interface tension $\gamma_{SE}$ on villus morphologies. A** Phase diagram showing simulation-end elongation length (circle colour, see heatmap) as a function of $\gamma_{SL}$ and $\gamma_{LE}$ when $\gamma_{SE} = 0.12$ (as opposed to $\gamma_{SE} = 0.06$ for all other simulations unless otherwise stated). The background colour of the phase diagram indicates the spreading regime, as indicated on top (blue is where S-tissue spreads ($\mathbf{S}_S > 0$), green is where L-tissue spreads ($\mathbf{S}_L > 0$), and orange is where E-tissue spreads ($\mathbf{S}_E > 0$)). Grey is the non-spreading regime ($\mathbf{S}_i < 0$ for $i = S, L, E$). **B,C** Snapshots of two elongating villi at consecutive time-points in a simulation, with each time-point shown below the snapshot. Snapshots are coloured by the concentration of $c_S$, $c_L$, and $c_E$ as indicated by the ternary plot with vertices of $c_S = c_L = c_E = 1$. $D = 1$, $\gamma_{SE} = 0.12$ and $\eta = 100$ for all simulations. See (A) for values of $\gamma_{LE}$ and $\gamma_{SL}$.
